# How robust are cross-population signatures of polygenic adaptation in humans?

**DOI:** 10.1101/2020.07.13.200030

**Authors:** Alba Refoyo-Martínez, Siyang Liu, Anja Moltke Jørgensen, Xin Jin, Anders Albrechtsen, Alicia R. Martin, Fernando Racimo

## Abstract

Over the past decade, summary statistics from genome-wide association studies (GWASs) have been used to detect and quantify polygenic adaptation in humans. Several studies have reported signatures of natural selection at sets of SNPs associated with complex traits, like height and body mass index. However, more recent studies suggest that some of these signals may be caused by biases from uncorrected population stratification in the GWAS data with which these tests are performed. Moreover, past studies have predominantly relied on SNP effect size estimates obtained from GWAS panels of European ancestries, which are known to be poor predictors of phenotypes in non-European populations. Here, we collated GWAS data from multiple anthropometric and metabolic traits that have been measured in more than one cohort around the world, including the UK Biobank, FINRISK, Chinese NIPT, Biobank Japan, APCDR and PAGE. We then evaluated how robust signals of polygenic score overdispersion (which have been interpreted as suggesting polygenic adaptation) are to the choice of GWAS cohort used to identify associated variants and their effect size estimates. We did so while using the same panel to obtain population allele frequencies (The 1000 Genomes Project). We observe many discrepancies across tests performed on the same phenotype and find that association studies performed using multiple different cohorts, like meta-analyses and mega-analyses, tend to produce polygenic scores with strong overdispersion across populations. This results in apparent signatures of polygenic adaptation which are not observed when using effect size estimates from biobank-based GWASs of homogeneous ancestries. Indeed, we were able to artificially create score overdispersion when taking the UK Biobank cohort and simulating a meta-analysis on multiple subsets of the cohort. Finally, we show that the amount of overdispersion in scores for educational attainment - a trait with strong social implications and high potential for misinterpretation - is also strongly dependent on the specific GWAS used to build them. This suggests that extreme caution should be taken in the execution and interpretation of future tests of polygenic score overdispersion based on population differentiation, especially when using summary statistics from a GWAS that combines multiple cohorts.

## Introduction

Most human phenotypes are polygenic: the genetic component of trait variation across individuals is caused by differences in genotypes at a large number of variants, each with a relatively small contribution to a given trait (Fisher et al., 1918; Turelli, 2017). This applies to phenotypes as diverse as a person’s height, their risk of schizophrenia or their risk of developing arthritis. The study of differences in complex traits spans more than a century but only in the last two decades has it become possible to systematically explore the underlying genetic architecture underlying these differences (Sella and N Barton, 2019). The advent of genome-wide association studies has led to the identification of thousands of variants that are associated with polygenic traits, either due to true biological mechanisms or because of linkage with causal variants (Visscher et al., 2012).

However, most research into the genetic aetiology of complex traits is based on GWAS data from populations of European ancestries (Popejoy and Fullerton, 2016). This bias in representation contributes to existing disparities in medical genetics and healthcare around the world (AR Martin, Kanai, et al., 2019). The low portability of European GWAS results - and, in particular, polygenic scores - to non-European populations is particularly concerning (AR Martin, Gignoux, et al., 2017; AR Martin, Kanai, et al., 2019) (but see Ragsdale et al., 2020). For example, the predictive accuracy of polygenic scores for height constructed using European effect size estimates has been shown to decrease with decreasing European ancestry in admixed populations (Bitarello and Mathieson, 2020). Recent studies have shown that ancestry deconvolution can be used to improve accuracy (Marnetto et al., 2020; M Wang et al., 2020), but important trait-associated variants in non-European populations may be missed if they have low frequencies or are absent in European populations. Moreover, effect size estimates for an associated variant derived from a European-ancestry GWAS may not accurately reflect the effect of the same variant on the trait in other populations (Wojcik et al., 2019). This could be due to differences in epistasis, differences in linkage disequilibrium between causal and ascertained variants, or gene-by-environment interactions, to name a few causes (Guo et al., 2018). Additionally, negative selection and demographic history may cause differences in genetic architectures between populations (Durvasula and Lohmueller, 2019).

During the last decade, GWAS summary statistics have also been used to look for evidence of directional selection pushing a trait to a new phenotypic optimum, via allele-frequency shifts occurring across a large number of associated variants - a phenomenon known as polygenic adaptation (Hayward and Sella, 2019; Pritchard et al., 2010). For example, several studies have consistently found evidence for polygenic adaptation operating on height-associated variants in Europe, mainly across a south-to-north gradient (Berg and Coop, 2014; Berg, Zhang, et al., 2017; Mathieson et al., 2015; Racimo et al., 2018; MR Robinson, Hemani, et al., 2015; Turchin et al., 2012). To test for selection, these studies primarily relied on summary statistics from the GIANT consortium dataset, which is a meta-analysis of anthropometric GWAS from multiple European cohorts (Allen et al., 2010; Wood et al., 2014). They looked for overdispersion and/or directional changes in the frequencies of trait-associated variants across populations, relative to a neutral null model. To account for potential confounding due to population stratification, some have tried to replicate this signal using family-based association studies (Allison et al., 1999; MR Robinson, Hemani, et al., 2015). Berg, Harpak, et al., 2019 and Sohail et al., 2019 showed that this signal of polygenic score overdispersion on height-associated variants in Europe (and possibly on other trait-associated variants) is attenuated and in some cases no longer significant when using effect size estimates from a GWAS performed on the UK Biobank – a large cohort composed primarily of individuals of British ancestry (Bycroft et al., 2018). There is no single explanation yet for these contradictory findings, but the most plausible one is that previous studies were impacted by very subtle confounding due to uncorrected population stratification in GIANT, and that data from family-based studies was not analyzed properly (Berg, Harpak, et al., 2019; Sohail et al., 2019).

It is as yet unclear how the choice of GWAS cohort affects tests of polygenic score overdispersion based on allele frequency differences between populations. Each cohort differs in ancestries of participants, inclusion criteria of individuals, SNP ascertainment scheme and association method. Given the poor portability of polygenic scores across populations, is it also true that GWASs performed on different cohorts will result in inconsistent signals of selection? Can we narrow down on the reason for the inconsistencies in previous studies of polygenic adaptation by looking at a larger number of cohorts? Here, we collated GWAS summary statistics from multiple complex traits that have been measured in more than one cohort around the world. We then evaluated how robust signals of polygenic score overdispersion are to the choice of cohort used to obtain effect size estimates. Across all comparisons, we used the same population genomic panel to obtain population allele frequency estimates: The 1000 Genomes Project phase 3 (The 1000 Genomes Project Consortium, 2015). We observe many discrepancies across tests performed on the same phenotype and attempt to understand what may be causing these discrepancies. We compare results for several traits and pay special attention to height, as it is the most well-characterized and studied complex trait in the human genetics literature, as well as a trait for which we have summary statistics from the largest number of GWAS cohorts. Finally, we perform an analogous analysis on educational attainment - a trait that has also been highlighted in recent studies of polygenic adaptation in humans (Racimo et al., 2018; Stern et al., 2020; Uricchio et al., 2019), and that is especially prone to be misinterpreted or misappropriated (Harmon, 2018; Novembre and NH Barton, 2018). We show that overdispersion signals for this trait are also highly sensitive to the choice of GWAS cohort.

## Methods

### GWAS summary statistics

We obtained GWAS summary statistics from five large-scale biobanks, a GWAS meta-analysis and a mega-analysis (Figure 1). Since we aim to make comparisons among them, our interest is focused on traits that were measured in at least two different cohorts. This resulted in a total of 30 traits being included in our analysis.

**Figure 1.**
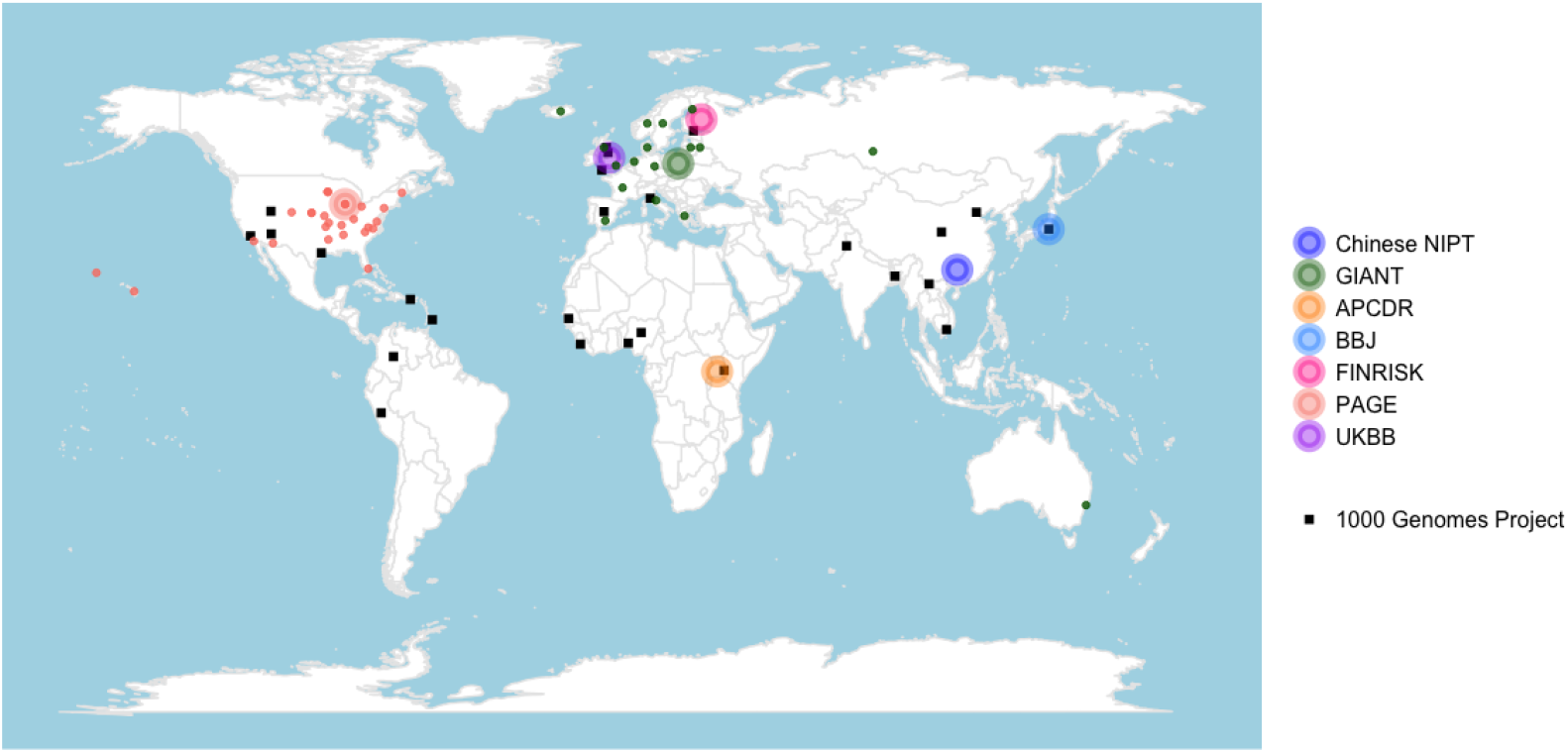
Map containing the geographic provenance of the panels of each association study we examined (large circles), as well as the 26 population panels from the 1000 Genomes Project (black squares). Small colored dots denote the provenance of the individual GWAS cohorts that were used in GIANT and in PAGE.

Below, we provide a brief summary of each of the GWASs we focused on. For an overview of the type of arrays and association methods used in each of these, see Table 1.

**Table 1.** Information on genotyping, imputation and association methods for each GWAS analyzed in this study. LM+PCs = linear model with principal components as covariates. LMM = linear mixed model.

| GWAS cohort | No. individuals | No. genotyped SNPs | No. imputed SNPs | Association method | GWAS array or sequencing method | Filters |
| --- | --- | --- | --- | --- | --- | --- |
| UKBB | 360388 | 820967 | 10,800,000 | LM+PCs | UK Biobank Axiom Array | INFO > 0.8<br>MAF > 1e-4<br>HWE P > 1e-10 |
| GIANT | 253288 | [meta-analysis] | 2,834,208 | LM+PCs | Affymetrix 500K platform, Illumina genotyping arrays and custom Perlegen arrays | Removed low quality SNPs |
| BBJ | 159,095 | 958,497 | 27,896,057 | LM+PCs | Illumina HumanOmniExpressExome | INFO ≥ 0.4<br>MAF > 1.0%<br>HWE P ≥ 1e-6 |
| Chinese NIPT | 61,321 | NA | 2,130,000 | LM+PCs | Ultra low depth shotgun sequencing | INFO > 0.4<br>HWE P > 1e-6 |
| PAGE | 49,781 | 1,748,250 | 39,723,562 | LMM | Illumina Multi-Ethnic Genotyping Array (MEGA) | INFO > 0.4 |
| FINRISK | 24,725 | 551,004 | 11,670,715 | LM+PCs | Illumina CoreExome | INFO > 0.7<br>MAF > 1.0%<br>HWE P > 1e-6 |
| APCDR | 4,778 | 2,382,209 | 16,477,797 | LMM | Illumina Omni2.5 | INFO ≥ 0.3<br>MAF > 0.5% |

- **UKBB**: Summary statistics from the GWAS performed on all UK Biobank traits (Bycroft et al., 2018). These were released by the Neale lab (round 2: http://www.nealelab.is/uk-biobank/), after filtering for individuals with European ancestries. The UK Biobank includes genetic and phenotypic data from participants from across the United Kingdom, aged between 40 and 69. The traits measured include a wide range of lifestyle factors, physical measurements, and other phenotypic information gained from blood, urine and saliva samples. The Neale lab performed association testing in ∼ 340,000 unrelated individuals.
- **FINRISK**: Summary statistics from GWASs carried out using the National FINRISK 1992-2012 collection from Finland (https://personal.broadinstitute.org/armartin/sumstats/finland/). The FINRISK study is coordinated by the National Institute for Health and Welfare (THL) in Finland and its target population is sampled from six different geographical areas in Northern Finland. The FINRISK cohort was conducted as a cross-sectional population survey every 5 years from 1972 to assess the risk factors of chronic diseases and health behavior in the working age population. Blood samples were collected from 1992 to 2012. Anthropometric measures and other lifestyle information were also collected. The number of samples used for the GWAS results varies among the different traits (∼25,000 to ∼5,000) (Borodulin et al., 2018).
- **PAGE**: Summary statistics from a multi-ethnic GWAS mega-analysis performed by the PAGE (Population Architecture using Genomics and Epidemiology) consortium (http://www.pagestudy.org/). This is a project developed by the National Human Genome Research Institute and the National Institute on Minority Health and Health Disparities in the US, to characterize population-level disease risks in various populations from the Americas (Carlson, 2016; Matise et al., 2011). The association analysis was assembled from four different cohorts: the Hispanic Community Health Study/Study of Latinos (HCHS/SOL), the Women’s Health Initiative (WHI), the Multiethnic Cohort (MEC) and the Icahn School of Medicine at Mount Sinai BioMe biobank in New York City (BioMe). The authors performed GWAS on 26 clinical and behavioural phenotypes. The study includes samples from 49,839 non-European-descent individuals. Genotyped individuals self-reported as Hispanic/Latino (n = 22,216), African American (n = 17,299), Asian (n = 4,680), Native Hawaiian (n = 3,940), Native American (n = 652) or Other (n = 1,052). The number of variants analyzed varies from 22 to 25 million for continuous phenotypes and 11 to 28 million for case/control traits. Sample sizes ranged from 9,066 to 49,796 individuals (Wojcik et al., 2019).
- **BBJ**: Summary statistics from GWASs performed using the Biobank Japan Project, which enrolled 200,000 patients from 12 medical institutions located throughout Japan between 2003-2008 (http://jenger.riken.jp/en/result). The authors collected biological samples and other clinical information related to 47 diseases and self-reported anthropometric measures. GWASs were then conducted on approximately 162,000 individuals to identify genetic variants associated with disease susceptibility and drug responses. Around 6 million variants were included for association testing (Hirata et al., 2017; Kanai et al., 2018; Nagai et al., 2017).
- **Chinese NIPT**: Summary statistics from a GWAS performed in China using non-invasive prenatal testing (NIPT) samples from ∼ 141,431 pregnant women (https://db.cngb.org/cmdb). The participants were recruited from 31 administrative divisions across the country. The study aimed to investigate genetic associations with maternal and infectious traits, as well as two antropometric traits: height and BMI (S Liu et al., 2018). It included ∼ 60,000 individuals. The number of imputed variants used was around 2 million.
- **APCDR**: Summary statistics performed using the African Partnership for Chronic Disease Research cohort, which was assembled to conduct epidemiological and genomic research of non-communicable diseases across sub-Saharan Africa (https://personal.broadinstitute.org/armartin/sumstats/apcdr/). The dataset includes 4,956 samples from Uganda (Baganda, Banyarwanda, Burundi, and others). The authors performed GWAS on 34 phenotypes, including anthropometric traits, blood factors, glycemic control, blood pressure, lipid tests, and liver function tests (Heckerman et al., 2016).
- **GIANT**: Summary statistics published by the Genetic Investigation of Anthropometric Traits consortium (2012-2015 version, before including UK Biobank individuals) (Locke et al., 2015; Wood et al., 2014)(https://portals.broadinstitute.org/collaboration/giant/index.php/GIANT_consortium_data_files). GIANT is a meta-analysis of summary association statistics for various anthropometric traits, and includes information from more than 250,000 individuals of European descent. The meta-analysis was performed on 2.5 million autosomal SNPs, after imputation.

### Population genomic panel

We used the 1000 Genomes Project phase 3 release data (The 1000 Genomes Project Consortium, 2015) to retrieve the allele frequencies of trait-associated variants in different population panels sampled from around the world (Figure 1). We used these to compute polygenic scores for each panel, using autosomal SNPs only. The dataset contains samples from 2,504 people from 26 present-day population panels, whose abbreviations and descriptions are listed in Table S1.

### Identifying trait-associated SNPs

We used summary statistics for a set of 30 traits that were measured in at least two of the previously-listed GWAS datasets. Table S2 shows the full list of the traits included in this analysis and the number of variants and individuals per trait. For each trait, we excluded triallelic variants, variants with a minor allele frequency lower than 0.01 across all samples and those classified as low confident variants whenever this information was available in the summary statistics file. We selected a set of trait-associated SNPs based on a P-value threshold, and the effect size estimates of these variants were used to construct a set of polygenic scores. To only include approximately independent trait-associated variants in our scores, we use a published set of 1,703 non-overlapping and approximately independent linkage-disequilibrium (LD) blocks to divide the genome (Berisa and Pickrell, 2016). We extracted the SNP within each block with the lowest association P-value. To investigate the robustness of signals to different filtering schemes, we used two P-value thresholds to extract significantly associated variants: 1) *P* < 1_e_−5 and 2) the standard genome-wide significant cutoff, *P* < 5_e_−8. Blocks that only contain variants that do not meet the chosen threshold were filtered out. As an example, Figure S1.A shows the distribution of effect size estimates of height-associated SNPs with *P* < 1_e_−5. In turn, Figure S1.B shows the distribution of the product of the effect size estimates and the square root of the study’s sample size (N). This serves as a fairer comparison among studies, as the standard error of the effect size estimate is approximately proportional to the inverse of 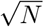 (see Casella and Berger, 2002; Edge, 2019; Holland et al., 2016). In order to build an empirical genome-wide covariance matrix (F-matrix) with non-associated SNPs, we extracted all SNPs with a P-value larger than 5_e_−8 and then sampled every 20th of these SNPs across the autosomal genome.

We also used the LD score regression approach (Heckerman et al., 2016) to obtain an LD score regression intercept, LD score regression ratio, and a SNP heritability estimate for each GWAS that we looked at. The LD score intercept is an estimate of the contribution of population stratification to test statistic inflation in a GWAS analysis. The LD score regression ratio measures the proportion of the inflation in the mean *χ*^2^ statistic that the LD Score regression intercept ascribes to causes other than polygenic heritability. Finally, an estimate of trait heritability can be obtained from the LD score regression slope (B Bulik-Sullivan et al., 2015; BK Bulik-Sullivan et al., 2015). We note, however, that Berg, Harpak, et al. (2019) showed that some of the assumptions of LD score regression - which allow one to separate estimates of stratification confounding from heritability - may be violated in the presence of background selection. Thus, these estimates may not accurately reflect the amount of stratification truly present in a GWAS.

### Neutrality test for polygenic scores

Polygenic risk scores are used to predict the genetic risk of a disease, or the genetic value of a trait, by combining the additive effect of a large number of trait-associated loci across the genome. For each trait, we obtained polygenic scores by computing the sum of allele frequencies at each of the top trait-associated SNPs from each block, weighted by their effect size estimates for that trait. The allele frequencies for these SNPs were retrieved from The 1000 Genomes Project population panels using *glactools* (Renaud, 2017). We then built a polygenic score vector for a given trait, 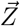, that contains the polygenic scores of all populations for that trait. Let 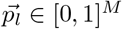 be the vector of derived allele frequencies at locus *l*, where *p*_*l,m*_ is the derived allele frequency at locus *l* in population *m*, while *α*_*l*_ is the effect size estimate of the derived allele at locus *l*. Then, the vector of the polygenic scores, 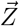, has length *M* equal to the number of populations (M = 26) and each element *Z*_*m*_ is the polygenic score for population *m*

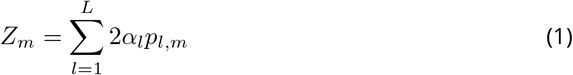

Here, *L* is the total number of trait-associated loci. For each polygenic score we built, we also obtained 95% credible intervals, constructed using the method in Sohail et al. (2019), assuming that the posterior distribution of the underlying population allele frequency is independent across populations and SNPs, and that it follows a beta distribution.

Berg and Coop, 2014 introduced a model designed for comparing polygenic scores across populations, in order to test for deviations from neutrality, which could perhaps be driven by adaptive divergence between populations. The test works by looking for overdispersion from a multivariate normal distribution, which would fit the distribution of scores if this was determined purely by genetic drift.

Under neutral genetic drift, Berg and Coop, 2014 showed that the joint distribution of 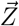 across closely-related populations should be approximately multivariate normal under a purely neutral model:

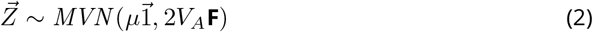

where 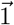 is a vector of ones and:

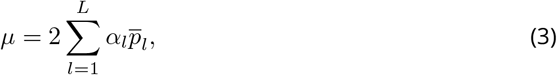

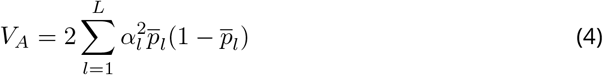

and 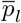 is the average allele frequency of locus *l* across all populations. The matrix **F** is a genome-wide covariance matrix that captures the co-ancestry among each pair of populations (Berg and Coop, 2014). Based on this null model, we can measure the Mahalanobis distance of the observed distribution of 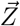 from the distribution under neutral genetic drift by computing *Q*_*x*_

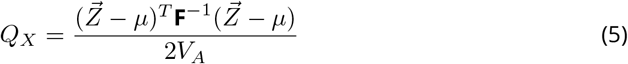

Under neutrality, the *Q*_*X*_ statistic is expected to follow a chi-squared distribution with M -1 degrees of freedom, 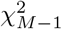 Berg and Coop, 2014). A significantly large value of *Q*_*X*_ indicates that there is an excess of variance in 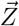 that cannot be explained by drift alone.

### P-values via randomization schemes

To avoid relying on the assumption that 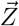 follows a multivariate normal distribution under neutrality, we also obtained P-values via two alternative methods (Berg and Coop, 2014; Berg, Zhang, et al., 2017; Racimo et al., 2018). The first one relies on obtaining neutral pseudo-samples by randomizing the sign (but not the magnitude) of the effect size estimates of all trait-associated SNPs, and then recomputing *Q*_*X*_. The second one involves obtaining pseudo-samples by sampling random SNPs across the genome with the same allele frequency distribution in a particular (target) population as the SNPs used to computed *Q*_*X*_. For each trait-associated SNP, we thus sampled a new SNP from a subset of the non-associated SNPs whose frequencies lie in the range [0.01 − *p, p* + 0.01] where *p* is the derived allele frequency of the trait-associated SNP. Then, we obtained a new P-value by computing the *Q*_*X*_ statistic on each of the pseudo-samples *i*:

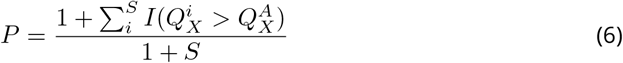

Here, 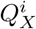 is the *Q*_*X*_ statistic computed on pseudo-sample *i*, 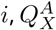 is the *Q*_*X*_ statistic computed on the true set of trait-associated SNPs, *I*() is an indicator function and *S* is the number of pseudo-samples used, which was set to 1,000. We tested the effect of using different population panels as our ‘target’ population for the frequency-matching scheme. Since we are utilizing seven GWAS cohorts that are composed of Latin American individuals, Asian (Japanese and Chinese), sub-Saharan African and European (Finnish and British) individuals, we decided to use population panels from the 1000 Genomes that roughly matched the ancestry of the GWAS cohorts: CHB for Chinese NIPT, JPT for BBJ, LWK for APCDR, FIN for FINRISK and GBR for UKBB. While PAGE is a very heterogeneous cohort, we find that PUR is the panel with the lowest amount of differentiation to PAGE, among all 1000G panels (Table S3), so we used PUR as the closest match to PAGE.

### Evaluating population structure

To look for population stratification along different axes of population variation Sohail et al., 2019, we first selected those variants that were present in the 1000 Genomes Project, the UKBB height GWAS and another non-UKBB height GWAS used for comparison against UKBB. We filtered out variants that had minor allele frequency < 5% in the 1000 Genomes Project, or that were located in the MHC locus (chr6:28477797-33448354) or in the chromosome eight inversion region (chr8:7643092-11528113). We then performed LD pruning on the resulting set of variants (using the –indep-pairwise 200 100 0.2 option). The remaining SNPs were used to perform a PCA on a matrix of genotypes from all the 1000 Genomes Project individuals, from which we obtained the first 20 PC loadings of population structure, using plink. Then, we performed linear regression of the PC scores on the genotypes of each SNP that was previously removed due to the pruning procedure. Finally, we plotted the correlations between each of the PCs and the effect size estimates from one of the two GWASs: UKBB or non-UKBB.

### Assessing different association methods

We were also interested in evaluating the effects of different types of association methods on the significance of the *Q*_*X*_ statistic. We used the UKBB cohort to perform different types of association studies on height. Starting from 805,426 genotyped variants across the genome, we restricted to SNPs with a minor allele frequency (MAF) > 5% globally, and performed associations on three different sets of individuals from the UKBB cohort: 1) self-reported white British individuals (“British”), 2) self-reported “white” individuals, and 3) “all ethnicities”, i.e. a UKBB set including all self-reported ethnicity categories. We applied the following quality filters in each of the resulting sets: 1) removed variants with *P* < 1_e_−10 from the Hardy-Weinberg equilibrium test, 2) removed variants with *MAF* < 0.1% in the set, 3) removed variants with an INFO score less than 0.8, 4) removed variants outside the autosomes, and 5) removed individuals that were 7 standard deviations away from the first six PCs in a PCA of the set (following the Neale lab’s procedure for defining British ancestry after performing PCA on the UKBB dataset). We then performed a GWAS via a linear model (LM) using PLINK v1.9 (Chang et al., 2015) and a GWAS via a linear mixed model (LMM) using BOLT-LMM (Loh et al., 2018), on each of the three sets (Table 2). We used sex, age, age^2^, sex*age, sex*age^2^ and the first 20 PCs as covariates.

**Table 2.** Description of types of methods used to obtain SNP associations from UKBB.

| Code | Cohort composition | Description of association method |
| --- | --- | --- |
| british linear-model | Self-reported "British" individuals from UKBB | linear model implemented in PLINK 1.9 |
| british mixed-model | Self-reported "British" individuals from UKBB | linear mixed model implemented in BOLT-LMM |
| white linear-model | Self-reported "White" individuals from UKBB | linear model implemented in PLINK 1.9 |
| white mixed-model | Self-reported "White" individuals from UKBB | linear mixed model implemented in BOLT-LMM |
| all ethnicities linear-model | All self-reported ethnicities from UKBB | linear model implemented in PLINK 1.9 |
| all ethnicities mixed-model | All self-reported ethnicities from UKBB | linear mixed model implemented in BOLT-LMM |
| meta-analysis.inverseSE | 75 sub-cohorts of equal size, composed of randomly sampled individuals from UKBB (all ethnicities) | inverse-variance-based meta-analysis implemented in METAL |
| meta-analysis.inverseSE | 75 sub-cohorts obtained from K-means clustering using 1st 3 PCs of UKBB individuals (all ethnicities) | inverse-variance-based meta-analysis implemented in METAL |
| meta-analysis.samplesize | 75 sub-cohorts of equal size, composed of randomly sampled individuals from UKBB (all ethnicities) | sample-size-based meta-analysis implemented in METAL |
| meta-analysis.samplesize | 75 sub-cohorts obtained from K-means clustering using 1st 3 PCs of UKBB individuals (all ethnicities) | sample-size-based meta-analysis implemented in METAL |

We also aimed to test whether a meta-analysis approach could lead to overdispersion of polygenic scores, and consequently, an inflated *Q*_*X*_ statistic. Therefore, we created a set of artificial meta-analyses on the entire UKBB cohort, approximately emulating the number of individual sub-cohorts that were included in GIANT. We used both the “all ethnicities” and the “white British” UKBB cohorts to compare the results of a meta-analysis on a homogeneous vs. a diverse cohort of individuals. For each of the two cohorts, we divided the corresponding set of individuals into 75 subsets, using two different approaches. In one approach, we obtained 75 clusters from a K-means clustering of the first three principal components from a PCA of the individuals. Under this approach, different cohorts have different sample sizes (though they do not exactly match the cohort size distribution observed in GIANT). In the other approach, we created 75 groups of equal size, randomly assigning individuals to each group, regardless of their placement in the PCA.

We used PLINK 1.9 to perform a linear association model in each of the 75 clusters or groups. As before, we used sex, age, age^2^, sex*age, sex*age^2^ and the first 20 PCs as covariates. These covariates were included in the analysis of each cohort before the meta-analysis. We explored how PC correction affected the meta-analyses. As the first 20 PCs, we used either the components from a PCA performed on each of the 75 sub-cohorts or the components from a PCA performed on all individuals together, before they were split. We note that the latter PCA would not be available to a researcher performing a meta-analysis in practice, but we carried it out to check whether lack of power to correctly model population structure via the cohort-specific PCAs was somehow misleading us. Afterwards, we integrated all summary statistics into a meta-analysis, using two different methods (Table 2): an inverse variance method and a sample size-based method, both implemented in METAL (Willer et al., 2010). This led to a total of 16 different types of meta-analyses artificially performed on the UKBB data.

### Educational attainment GWAS

We also performed an assessment of robustness in the distribution of population-wide polygenic scores for educational attainment. In this case, together with effect size estimates from the UK Biobank, we also obtained estimates from three studies carried out by the Social Science Genetic Association Consortium (SSGAC) (https://www.thessgac.org):

- A meta-analysis of 126,559 individuals (42 discovery cohorts and 12 replication cohorts) (Rietveld et al., 2013)
- A meta-analysis of 293,723 individuals (64 cohorts) (Okbay et al., 2016).
- A meta-analysis of 1,131,881 individuals (Lee et al., 2018) (71 cohorts in total). Note that this study includes the samples from the Okbay et al., 2016 study and the UK Biobank as well.

We took the summary statistics of each educational attainment GWAS “as is”, without modeling sample overlap between cohorts.

## Results

### Robustness of signal of selection and population-level differences

We obtained sets of trait-associated SNPs for GWASs performed on seven different cohorts: UK Biobank, FINRISK, Chinese NIPT, Biobank Japan, APCDR and PAGE. Using the effect size estimates from each GWAS, we calculated population-wide polygenic scores for each of the 26 population panels from the 1000 Genome Project (The 1000 Genomes Project Consortium, 2015), using allele frequencies from each population panel. We then tested for overdispersion of these scores using the *Q*_*X*_ statistic, which was designed to detect deviations from neutral genetic drift affecting a set of trait-associated SNPs (Berg and Coop, 2014). We focused on 30 traits that were phenotyped in two or more cohorts, so that we could compare the P-value of this statistic using effect size estimates from at least two different cohorts (see Methods).

We applied the *Q*_*X*_ statistic to each of the 30 traits by selecting SNPs we deemed to be significantly associated with each trait. We used two different P-value cutoffs to select these SNPs: 1) a lenient cutoff, *P* < 1_e_−5 and 2) the standard genome-wide significance cutoff *P* < 5_e_−8. To verify that significant P-values of the *Q*_*X*_ statistics were not due to violations of the chi-squared distributional assumption, we also computed P-values using two randomization schemes: one is based on randomizing the sign of the effect size estimates of the trait-associated SNPs (but not their magnitude), while the other is based on using frequency-matched non-associated SNPs (see Methods). In general, P-values obtained from the three schemes are broadly similar across the various approaches used. However, we observe a few inconsistencies in the sign-randomization scheme, when compared to the other two approaches (Figures S2 and S3). The number of significant SNPs for each of the traits under the two cutoffs is shown in Figures S4 and S5.

We used two types of multiple-testing Bonferroni corrections: one that applies to us - correcting for both the number of traits assessed and the number of cohorts on which each of those traits were tested (we call this number *m*) - and another that would apply to a person that was blind to the other cohorts - and so would only correct for the *n* traits tested within their available cohort (Figures 2, S2). We only find few traits with significant overdispersion in *Q*_*X*_. Under the *P* < 1_e_−5 SNP-association cutoff, the only traits with significant overdispersion in at least one cohort are height and white blood cells (WBC) (Figures 2, S2). Potassium levels in urine and mean corpuscular hemoglobin (MCH) also result in significant values of *Q*_*X*_ when using the *P* < 5_e_−8 SNP-association cutoff (Figures S3, S6).

**Figure 2.**
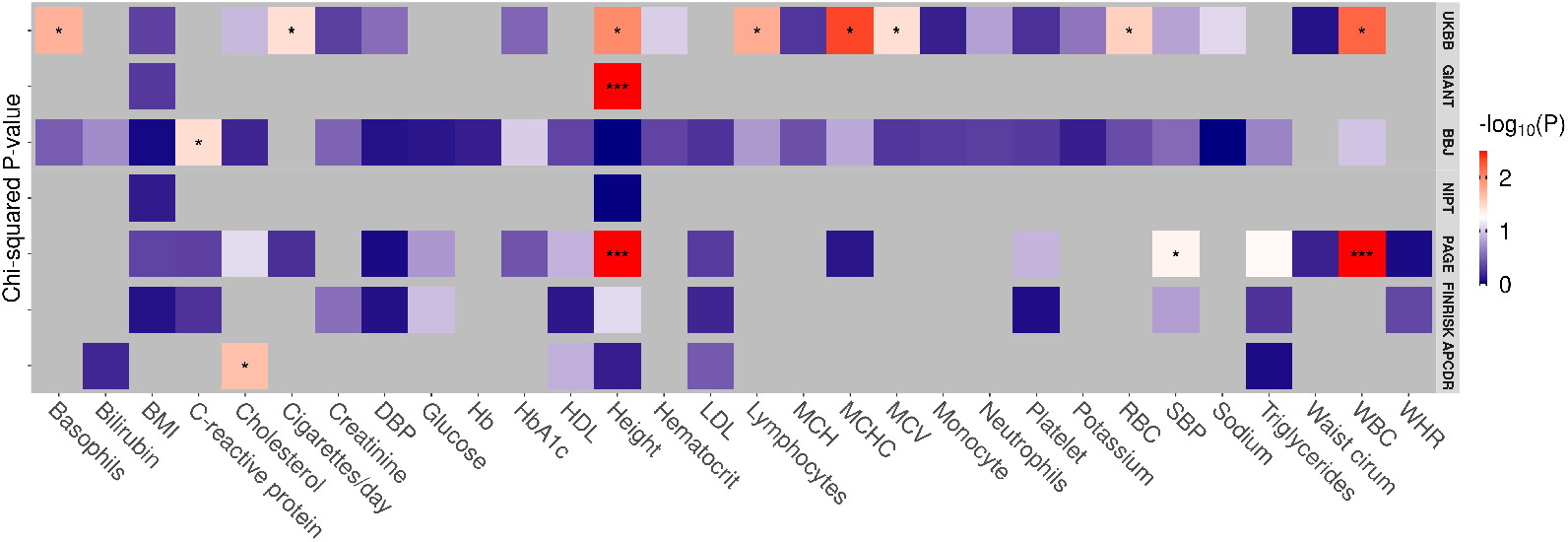
Each cell of this heatmap is the −*log*_10_(P-value) of a *Q*_*X*_ test using SNPs associated with different traits and effect size estimates obtained from different GWASs. Each row corresponds to one of the six GWASs we are evaluating and the columns correspond to the different traits for each GWAS that have SNPs with *P* < 1_e_−5. BMI, body mass index; DBP, Diastolic blood pressure; HbA1c, glycated hemoglobin; HDL, high-density lipoprotein; LDL, low-density lipoprotein; MCH, mean corpuscular hemoglobin; MCHC, mean corpuscular hemoglobin concentration; MCV, mean corpuscular volume; RBC, red blood cell count; SBP, systolic blood pressure; WBC, white blood cell count; WHR, waist-to-hip ratio. Significance thresholds after Bonferroni correction: *** < 0.05/m, ** < 0.05/n, * < 0.05, where *n* is the number of traits in each GWAS (row-dependent) and *m* is the total number of tests calculated, across all GWASs.

Figure 3 shows polygenic scores computed for each of the 1000 Genomes populations for height. In agreement with previous studies (Berg, Harpak, et al., 2019; Sohail et al., 2019), we observe that differences in polygenic height scores when using effect size estimates from the UKBB are greatly attenuated relative to differences in scores built when using estimates from GIANT. Extending this analysis across all datasets, we observe that PAGE polygenic scores are also over-dispersed, though in different directions than GIANT scores (Figure 3). Additionally, the observation that Europeans have very high polygenic scores when using GIANT effect size estimates cannot be replicated using any of the other GWAS estimates. After multiple testing correction (for both association P-value threshold schemes), we only obtain significant *Q*_*X*_ P-values when using summary statistics from PAGE and GIANT. The number of SNPs used for polygenic scores are shown in Table 3. The LD score regression ratio is substantially higher for PAGE and FINRISK than for the other cohorts (Tables 4, S4).

**Table 3.** Height *Q*_*X*_ scores and P-values, assuming a chi-squared distribution for the scores. The number of trait-associated SNPs used to compute the scores are shown for both cutoffs.

| GWAS cohort | Lenient association threshold ($P < 1e-5$) | | Strict association threshold ($P < 5e-8$) | |
| --- | --- | --- | --- | --- |
| | Num. trait associated SNPs | $Q_X$ | Num. trait associated SNPs | $Q_X$ |
| UKBB | 1140 | 45.57 ($P = 0.01$) | 810 | 39.72 ($P = 0.04$) |
| GIANT | 739 | 102.73 ($P = 4e-3$) | 468 | 69.18 ($P = 8e-8$) |
| BBJ | 759 | 9.255 ($P = 0.99$) | 431 | 35.00 ($P = 0.11$) |
| Chinese NIPT | 164 | 12.82 ($P = 0.99$) | 60 | 24.68 ($P = 0.54$) |
| PAGE | 528 | 77.31 ($P = 5e-7$) | 84 | 49.52 ($P = 4e-3$) |
| FINRISK | 209 | 36.43 ($P = 0.08$) | 47 | 41.48 ($P = 0.03$) |
| APCDR | 61 | 21.15 ($P = 0.73$) | 0 | NA |

**Table 4.** SNP-based heritability and LD Score regression ratio and intercept estimates (with standard errors in parentheses) for height measured in different cohorts. LD scores were computed for the super-population panels in the 1000 Genomes Project. The APCDR heritability estimate is not shown because it was estimated to be negative, due to the small sample size of the cohort.

| GWAS cohort | Genome-wide significant SNPs ($P < 5e-8$) | Observed scale heritability (SE) | LD regression ratio (SE) | LD regression intercept (SE) | Super Population |
| --- | --- | --- | --- | --- | --- |
| UKBB | 30891 | 0.4205 (0.0183) | 0.1184 (0.0098) | 1.4384 (0.0362) | EUR |
| GIANT | 11625 | 0.3159 (0.0146) | 0.1419 (0.0094) | 1.2727 (0.0181) | EUR |
| BBJ | 9976 | 0.4168 (0.019) | 0.1142 (0.0128) | 1.1829 (0.0204) | EAS |
| Chinese NIPT | 1573 | 0.329 (0.0201) | 0.0746 (0.0353) | 1.0425 (0.0201) | EAS |
| PAGE | 564 | 0.2578 (0.0237) | 0.371 (0.0336) | 1.1762 (0.016) | AMR |
| FINRISK | 415 | 0.4557 (0.0446) | 0.3775 (0.035) | 1.1358 (0.0126) | EUR |
| APCDR | 0 | NA | NA | NA | AFR |

**Figure 3.**
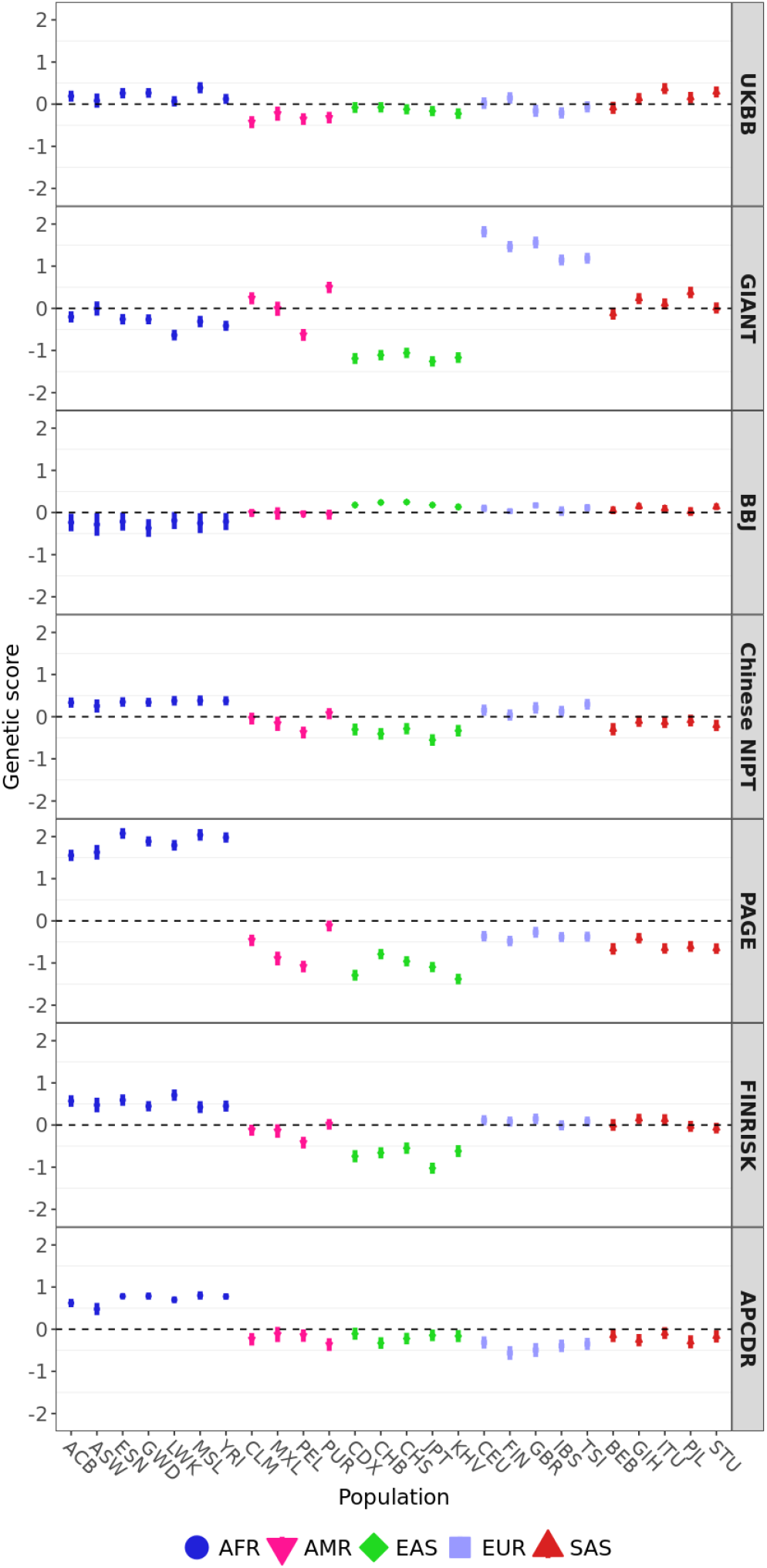
Polygenic scores for height using candidate SNPs with *P* < 1_e_−5. The 1000 Genomes Project populations colored by their super-population code. The corresponding number of trait-associated SNPs and the *Q*_*X*_ P-value for each GWAS are shown in Table 3. Error bars denote 95% credible intervals, constructed using the method in Sohail et al., 2019, assuming that the posterior distribution of the underlying population allele frequency is independent across populations and SNPs.

We also tested how the choice of the SNP association P-value threshold influenced the results. Sohail et al., 2019 showed that between-population differences in polygenic height scores grow stronger when using more lenient SNP association P-value cutoffs. However, one then runs the risk of including more variants that may be significantly associated due to uncorrected population stratification. We see there is a smaller score overdispersion when using the genome-wide significant SNPs, than when using the more lenient P-value cutoff (right column, Table 3 and Figure S7).

Finally, we computed polygenic scores on a single set of candidate SNPs ascertained in the largest biobank (UKBB) but using effect size estimates from each of the other GWAS in turn. Signals of overdispersion in height polygenic scores are greatly attenuated in each of the non-UKBB GWAS, and are much more similar to the patterns observed in UKBB (Figure S8 and Figure S9). This suggests an important reason for the observed overdispersion patterns in these other GWAS is the choice of significant SNPs recovered from each study.

We also looked in closer detail at other traits with evidence for significant overdispersion via the *Q*_*X*_ test. White blood cell counts (WBC), for example, shows strong overdispersion when using PAGE, but not when using the UKBB or BBJ effect size estimates (Figure S10). We also observe a similar pattern when looking at mean corpuscular hemoglobin (MCH) scores (Figure S11). In the case of potassium levels in urine, larger between-population differences are found in UKBB than in BBJ, when we use the stringent threshold (Figures S12). In general, we observe that between-population differences in polygenic scores tend to be more similar between studies when using the stricter SNP-association P-value threshold, than when using the more lenient threshold.

### Relationship between GWAS effect size estimates

To better understand where the differences in overdispersion of *Q*_*X*_ could stem from, we performed pairwise comparisons of the effect size estimates from the different GWAS. Since the UKBB GWAS is the GWAS with the largest number of individuals, we decided to compare the estimates from each of the other studies to the UKBB estimates. Here, we only focused on the 1,703 approximately-independent SNPs (the best tag SNP within each LD block). We began by only using SNPs that were classified as significant in UKBB using the lenient cutoff (*P* < 1_e_−5) (Figure 4). We observe that effect size estimates are correlated, as expected, but the strength of this correlation varies strongly across comparisons. UKBB vs. GIANT shows the highest correlation, while UKBB vs. APCDR shows the lowest. Those SNPs that also have a significant P-value in the non-UKBB GWAS in each comparison (colored in red in Figure 4) show a higher correlation than the rest of the SNPs: a pattern expected due to the winner’s curse, and exacerbated by differences in sample sizes and LD patterns between GWAS cohorts (Berg, Harpak, et al., 2019).

**Figure 4.**
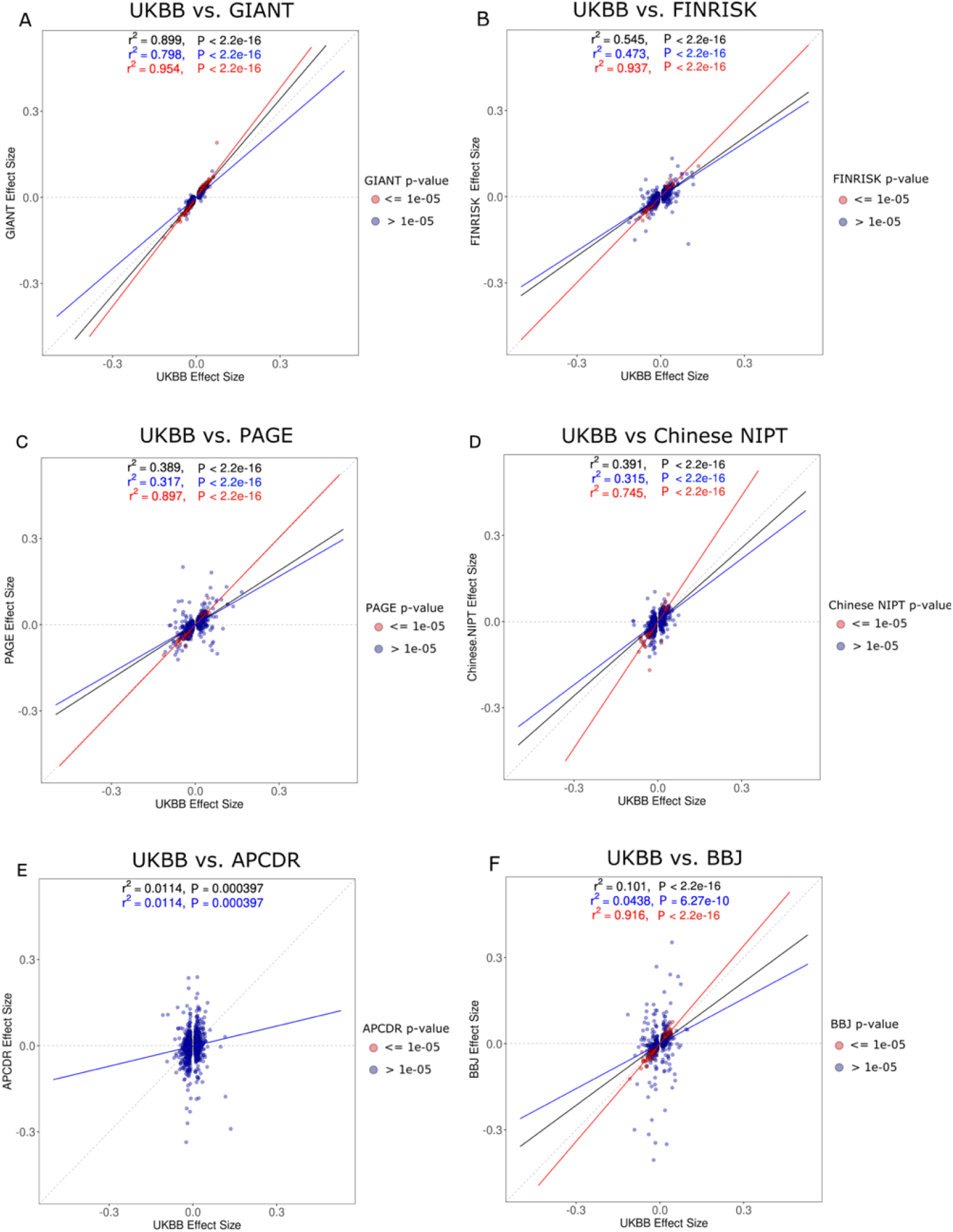
Regression of UKBB effect size estimates against non-UKBB effect size estimates, after filtering for SNPs with *P* < 1_e_−5 in the UKBB GWAS. The SNPs are colored based on their P-value in the non-UKBB GWAS, as are the corresponding regression lines (red: *P* < 1_e_−5; blue: *P* > 1_e_−5). The black regression line was obtained using all SNPs, regardless of their P-value in the non-UKBB GWAS.

The same analysis was carried out with SNPs classified as significant in each of the non-UKBB studies. The correlation of effect size estimates is generally lower (Figure S13), and a high percentage of SNPs deemed to be significant in the non-UKBB GWAS have effect size estimates approximately equal to zero in the UKBB GWAS (Figure S13). This pattern is stronger when we do not filter the 1,703 approximately independent SNPs by a particular SNP-association P-value cutoff (Figures S14 and S15).

We computed pairwise Pearson correlation coefficients between estimated effect sizes in the UKBB GWAS and each of the other GWAS (Table S5 when using SNPs that are significant in UKBB and Table S6 when using SNPs that are significant in the other GWAS). We observe that GWAS performed on individuals living geographically close to Britain have higher correlations to UKBB estimates than those that are performed on distant individuals. For instance, GIANT and FINRISK (both European-based GWAS), show high correlation in effect size estimates with UKBB (0.9958 and 0.790, respectively). In contrast, the GWAS carried out on an African cohort - APCDR - shows an extremely low correlation in effect size estimates with UKBB (correlation coefficient = 0.087). This cohort has by far the smallest sample size of all the cohorts we analyzed (n = 4,778), which may explain the low correlation. We also observe higher correlations when filtering for significantly associated SNPs using either of the two SNP significance thresholds (*P* < 1_e_−5 and *P* < 5_e_−8) from the 1703 LD blocks.

The sample size of the GWAS might also affect the correlation in effect size estimates. We can see in Figure S16 that there is a positive relationship between the *log*_10_ of the number of samples included in the non-UKBB GWAS and the Pearson correlation coefficients between the estimated effects in the non-UKBB GWAS and those estimated in the UKBB GWAS (which has the largest sample size). We note, however, that the slope of a linear regression between these variables is only significantly different from zero when ascertaining SNPs in the non-UKBB study, and filtering for SNPs with P < 1_e_−5. Most of our results stem from using LD block partitions derived from a European panel of the 1000 Genomes Project (Berisa and Pickrell, 2016; The 1000 Genomes Project Consortium, 2012). To investigate the sensitivity of our results to the choice of LD blocks (particularly when querying non-European GWASs), we also show results under an LD blocking scheme obtained from a population panel that was close to the GWAS from which we obtained effect size estimates (Berisa and Pickrell, 2016). For BBJ and Chinese NIPT, we use a set of 1,445 LD blocks constructed using LD patterns in the East Asian panels of the 1000 Genomes Project. For APCDR, we use a set of 2,582 LD blocks constructed using LD patterns in the African panels of the 1000 Genomes Project (Table S7). In the case of PAGE, we do not have a Latin-American-specific LD block partitioning scheme. However, when we use European LD blocks, we can detect a significant residual population structure pattern in PAGE, but this pattern is no longer significant when we use African LD blocks (Figure S17), so we show PAGE results using African LD blocks in Table S7.

### Evidence for population stratification

Berg, Harpak, et al., 2019 looked for latent population stratification by studying the relation between allele frequency differences in two populations and the difference in effect size estimates in two GWAS. Presumably, if neither GWAS is affected by population stratification, there should not be a relation between these two variables. We plotted SNP differences in allele frequency between northern European and East Asian, African, and southern European samples (GBR, CHB, LWK and TSI subsets of 1000 Genomes, respectively) against the difference in height effect size between a pair of GWAS. When comparing the UKBB and GIANT, we replicate the signal of correlation in differences between northern and southern European from Berg, Harpak, et al., 2019 (*P* < 1_e_−5, see Figure S18). This pattern is also observed in the GBR vs. CHB and GBR vs. LWK comparisons (Figures S19 and S20, panel A and B). However, these differences are not observed for any other pairwise GWAS comparisons (Figures S18, S22 and S23). We can see SNPs with large effect size differences tend to be low-frequency SNPs, as the standard error of the effect size estimate for a SNP in a GWAS is a function of the SNP’s expected heterozygosity (Holland et al., 2016).

We also followed Sohail et al., 2019’s approach to look for GWAS stratification along different PCA axes of population structure. We first performed a PCA on a matrix of genotypes from all 1000 Genomes Project individuals. Then, we computed the correlation between the first 20 loadings of that PCA and the effect size estimates obtained from the UKBB height GWAS, as well as the correlation between the same PC loadings and the effect size estimates from a different (non-UKBB) height GWAS, on the same set of sites (see Methods). We plotted each of these PC-specific correlations and colored them by the correlation between the corresponding PC and the allele frequency differences between two population panels: GBR vs. TSI (Figure 5); and GBR vs. CHB (Figure S24). This allows us to compare patterns of stratification between two GWASs (UKBB and non-UKBB) along particular axes of genetic variation. We observe large correlations between axes of population structure and effect size estimates in GIANT and PAGE, and to a lesser degree in FINRISK, but not in the other GWAS we queried. Overall, this suggests GIANT and PAGE might be more strongly confounded by stratification than the other GWASs under study.

**Figure 5.**
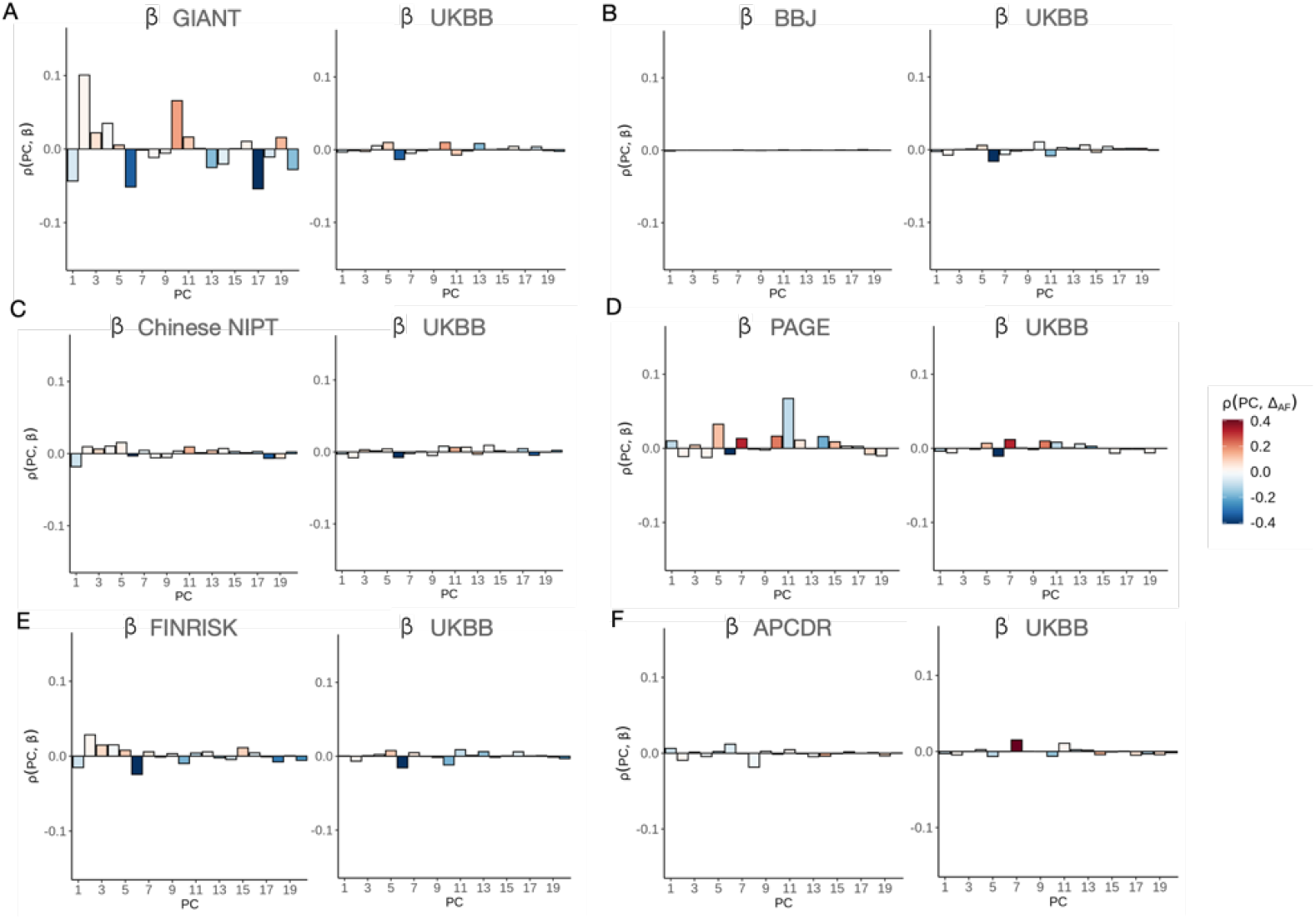
Pearson correlations between 20 PC loadings and height effect size estimates from a non-UKBB GWASs, compared to the same correlation using effect size estimates from the UKBB GWAS, for different choices of non-UKBB GWAS. The correlations were computed using SNPs that are present in both the UKBB and non-UKBB GWAS cohorts, and in the 1000 Genomes Project. The barplots are coloured by the correlation between each loading and the allele frequency difference between GBR and TSI. A) GIANT vs. UKBB. B) BBJ vs. UKBB. C) Chinese NIPT vs. UKBB. D) PAGE vs. UKBB. E) FINRISK vs. UKBB. F) APCDR vs. UKBB.

### Assessing different association methods

We find strong differences in the amount of polygenic score overdispersion across GWASs, but the GWASs we assessed were carried out using different association methods. We wanted to evaluate the effect of different association methods on the overdispersion of polygenic scores, while using the same underlying association cohort. We chose the UKBB cohort for this assessment, as it is the largest cohort among the ones we tested. We first split the UKBB cohort into three increasingly more expansive sets: 1) “British”, 2) “White”, and 3) “all ethnicities”, based on a self-identified ethnicity classification carried out by the UKBB consortium. We then performed linear model (LM) and linear mixed model (LMM) association methods on each of the three sets of individuals (Table 2, see Methods). We also wanted to see if we could replicate the strong overdispersion in polygenic scores we saw in GIANT, by partitioning the entire UKBB cohort into 75 cohorts (approximately emulating the number of cohorts in GIANT), and then performing a meta-analysis on the summary statistics obtained from individual GWASs performed separately on each of these cohorts (Table 2, see Methods).

The population-wide polygenic scores and the *Q*_*X*_ scores obtained using effect size estimates from each of these different methods are in Figures 6 and 7, respectively. We have more power to detect height-associated SNPs when we used the mixed model, and the distribution of polygenic scores differs quite markedly between the linear model and the mixed model approaches. For example, African polygenic scores tend to be higher when using the mixed model approach. Regardless of whether one uses a linear or a mixed model, GWASs performed on a more expansive category of people (“all-ethnicities”) lead to increased overdispersion of polygenic scores than when using more restrictive categories (“British” or “White”). Additionally, our artificial meta-analysis on the UKBB data resulted in even stronger overdispersion of the scores and, consequently, an even more strongly inflated *Q*_*X*_ statistic, regardless of the set used. However, polygenic scores dispersion is slightly attenuated when we use the “British” set. The most extreme *Q*_*X*_-derived P-values across settings were those from the meta-analyses in the “all ethnicities” set (Figure S25). This increased overdispersion is particularly evident when looking at the scores of European (FIN, CEU, GBR) and Latin American (PEL, CLM, MXL) populations in the meta-analysis setting (Figure 6). The choice of method for PC correction against population stratification does not lead to notably different results (Figure S26).

**Figure 6.**
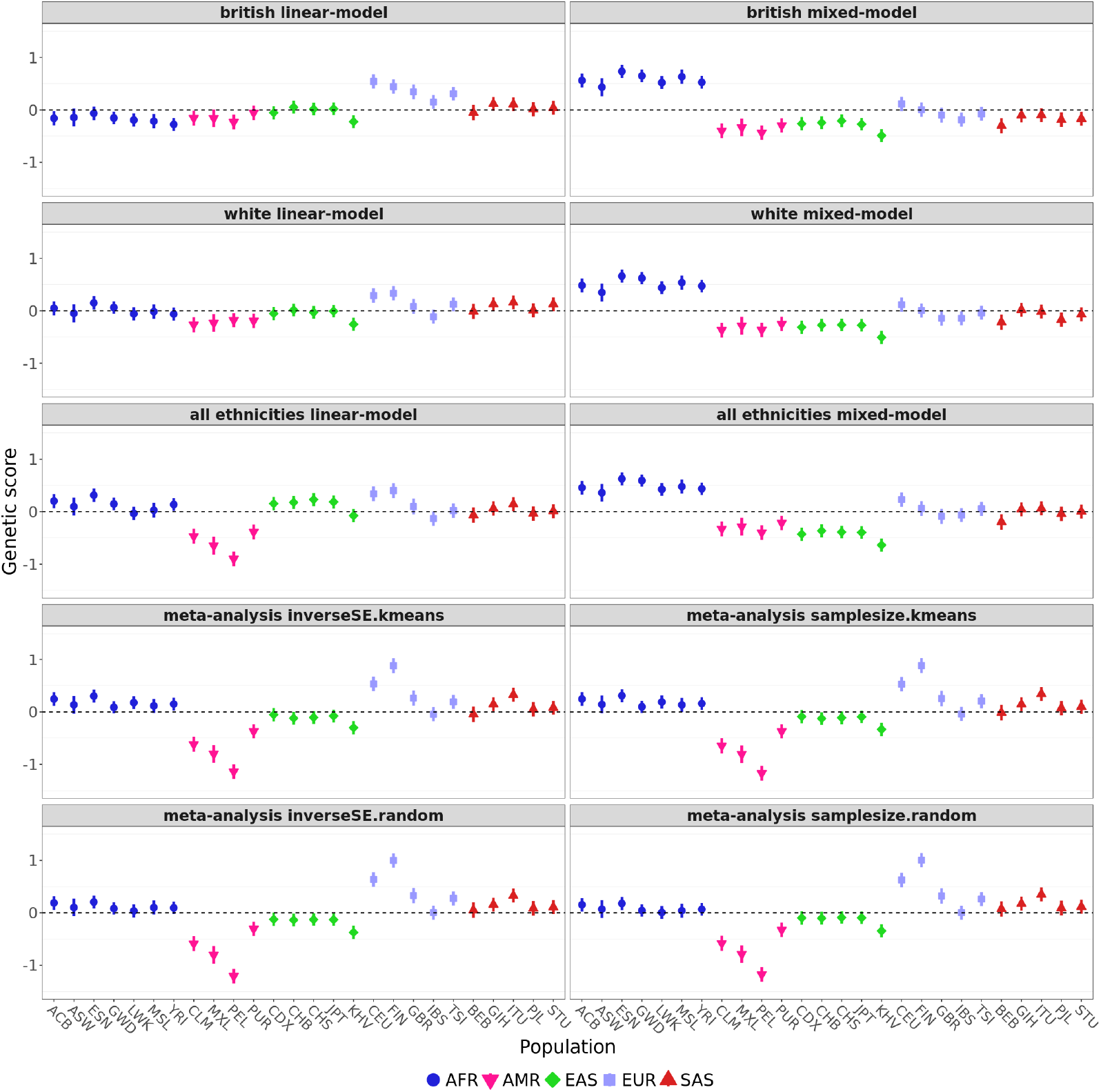
Polygenic scores for height in each of the 1000 Genomes population panels, using effect size estimates from UKBB, obtained via different types of association methods. The types of GWAS methods used are described in Table 2. Error bars denote 95% credible intervals, constructed using the method in Sohail et al., 2019, assuming that the posterior distribution of the underlying population allele frequency is independent across populations and SNPs.

**Figure 7.**
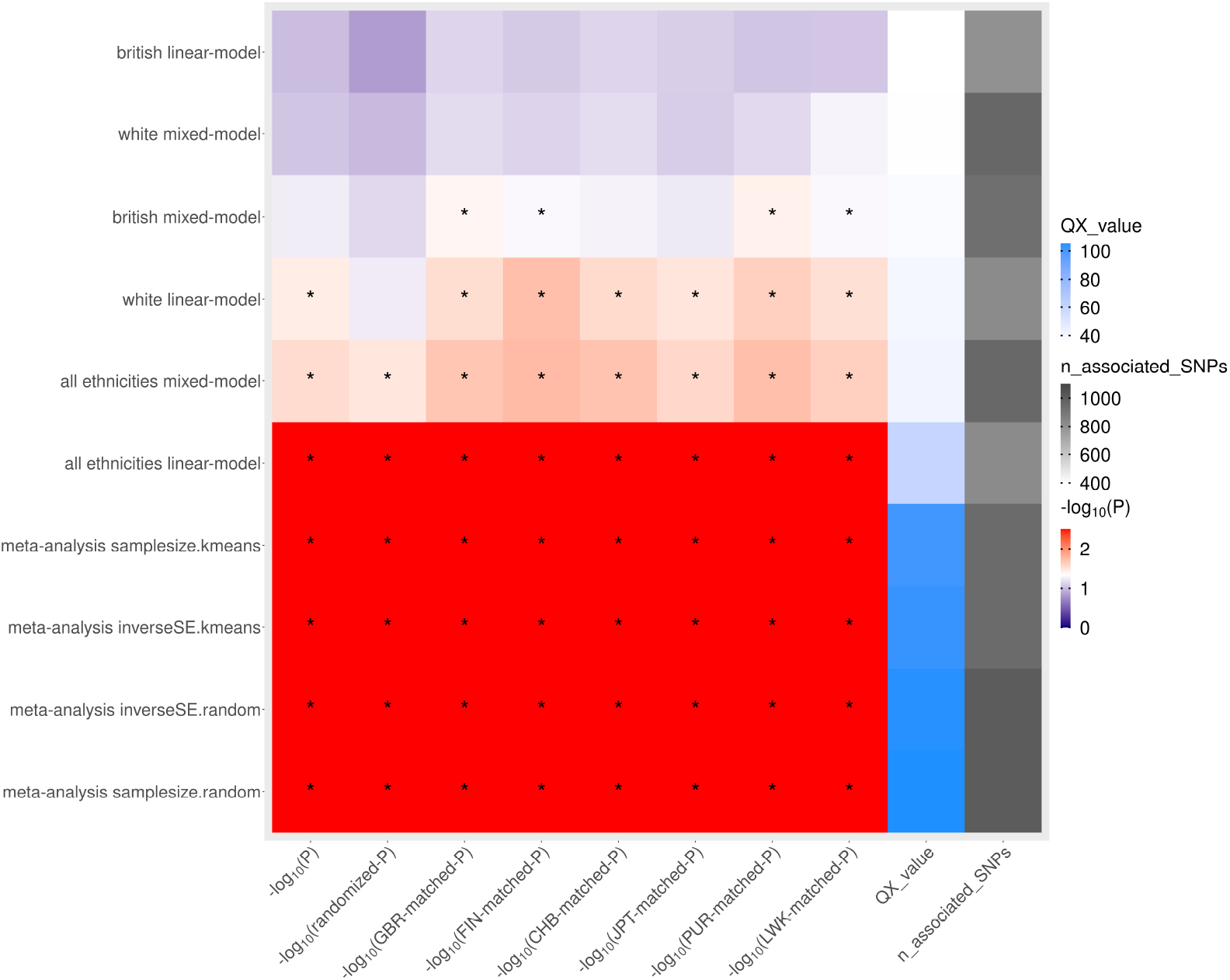
*Q*_*X*_ statistics and P-values for the trait height, obtained by using different types of association methods on the UKBB data. The asterisk denotes a significance cutoff for *Q*_*X*_ of *P* < 0.05. “-log10(Chi squared P)” = -log10(P), obtained assuming a chi-squared distribution for the *Q*_*X*_ statistic. “-log10(randomized P)” = -log10(P-value), obtained using the effect sign-randomization scheme. All other P-values were obtained by sampling random SNPs from the genome using the allele frequency matching scheme in different populations, as described in the Methods section. The different types of GWAS methods along the y-axis are described in Table 2.

### The case of educational attainment

Previous studies have also found evidence for strong score overdispersion in variants associated with educational attainment (Racimo et al., 2018; Uricchio et al., 2019) - a trait consisting in the number of school years an individual has received. This trait has received considerable attention in both the media and the scientific literature, due to its potential for misappropriation and misuse by far-right groups (Harmon, 2018; Novembre and NH Barton, 2018). Though associated variants have been shown to be disproportionally located in genes involved in brain development, this trait is also highly affected by the environment (Lee et al., 2018; Okbay et al., 2016), and potentially likely to be confounded by unaccounted factors, such as cultural and socioeconomic background, and parental salary and education. Like height, the genetic value of this trait has also been shown to be significantly similar among spouses, due to assortative mating (Abdellaoui et al., 2015; Hugh-Jones et al., 2016; MR Robinson, Kleinman, et al., 2017; Yengo et al., 2018), which could, in turn, affect the interpretation of population genetic tests that assume individuals are not segregating on the basis of trait preferences. The robustness of previously reported signals of overdispersion thus warrants some attention, and we therefore set out to assess how consistent patterns of overdispersion were across available GWAS cohorts.

Though educational attainment is only available in one of the cohorts we had access to (UKBB), there are also 3 publicly available GWAS on this trait that were carried out in individuals of European ancestry by the SSGAC consortium: Lee et al. (2018), Okbay et al. (2016), and Rietveld et al. (2013). The SSGAC consortium used increasingly larger meta-analyses to test for genetic associations with this trait (Lee et al., 2018 included both the Okbay et al., 2016 cohorts and the UK Biobank cohort in its meta-analysis). To be able to replicate the results of Racimo et al., 2018, we also computed polygenic scores on the Okbay et al., 2016 GWAS estimates using the posterior-probability approach (PPA) used in that study, and compared them to the P-value approach used throughout this manuscript.

As with height, we find strong inconsistencies in patterns of score dispersion, depending on the P-value cutoff used to include SNPs in the polygenic score, and on which cohort we use for deriving effect size estimates (Figure 8 and Table 5). For example, when using genome-wide significant SNPs to build scores, the strongest pattern of overdispersion is found when utilizing estimates from Okbay et al., 2016 with the PPA method (*Q*_*X*_-derived *P* = 0.0037). When including SNPs into the score via the more lenient SNP association P-value cutoff (1_e_−5), the strongest pattern of ovedispersion is found when using estimates from Lee et al., 2018 (*Q*_*X*_-derived *P* = 1.397_e_−5). Importantly though, the patterns of dispersion are different under these two conditions: European polygenic scores are highest in the latter, but East Asian scores are highest in the former. The UKBB score pattern when using genome-wide significant SNPs resembles the Okbay et al., 2016 pattern (as noted in Racimo et al., 2018) but is very different from the Lee et al., 2018 pattern, and is also different from scores derived from the same (UKBB) cohort under the more lenient association P-value scheme. Even though evidence for population stratification is weaker in these meta-analyses than in the height GWAS meta- and mega-analyses (Figure S27), the axes of stratification still differ across studies, which might help to explain the differences in score distributions.

**Table 5.** Educational attainment *Q*_*X*_ scores, assuming a chi-squared distribution for the scores. The number of trait-associated SNPs used to compute the scores are shown for both SNP P-value cutoffs.

| GWAS cohort | Lenient association threshold ($P < 1e-5$) | | Strict association threshold ($P < 5e-8$) | |
| --- | --- | --- | --- | --- |
| | Num. trait associated SNPs | $Q_X$ | Num. trait associated SNPs | $Q_X$ |
| UKBB | 746 | 40.087 ($P = 0.038$) | 246 | 42.859 ($P = 0.02$) |
| Leeetal.2018 | 907 | 67.761 ($P = 1.4e-5$) | 416 | 51.62 ($P = 0.002$) |
| Okbayetal.2016 | 319 | 36.814 ($P = 0.08$) | 74 | 35.175 ($P = 0.1$) |
| Rietveldetal.2013 | 54 | 35.444 ($P = 0.102$) | 4 | 29.005 ($P = 0.311$) |

**Figure 8.**
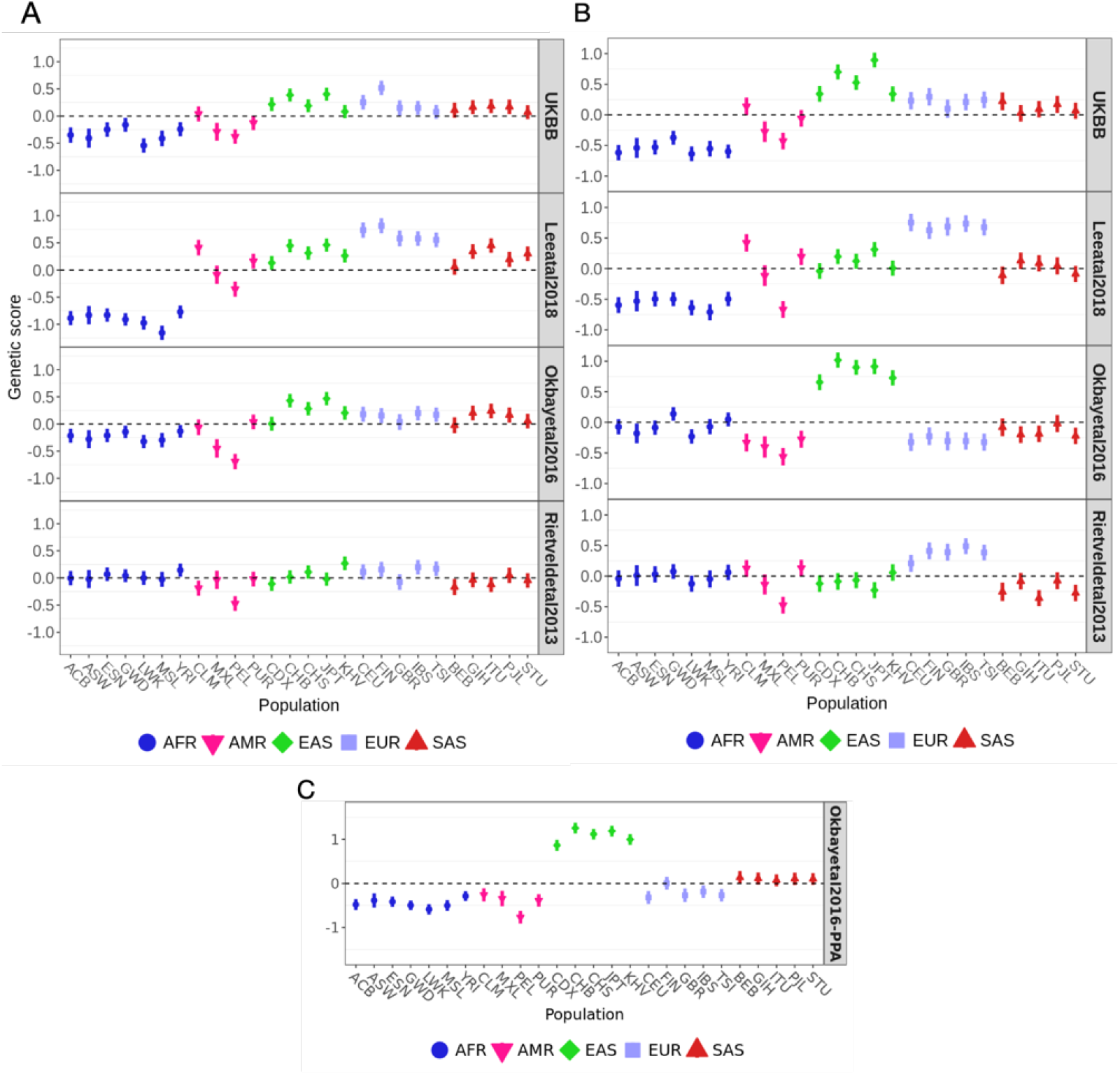
Polygenic scores for educational attainment in each of the 1000 Genomes population panels, using effect size estimates from three SSGAC consortium GWAS studies (Lee et al., 2018; Okbay et al., 2016; Rietveld et al., 2013) and the UKBB Neale lab effect size estimates. A. Polygenic scores for variants with *P* < 1_e_−5. B. Polygenic scores for variants with *P* < 5_e_−8. C. Variants selection using a posterior-probability approach (PPA) on the Okbay et al., 2016 summary statistics, emulating Racimo et al., 2018. Error bars denote 95% credible intervals, constructed using the method in Sohail et al., 2019, assuming that the posterior distribution of the underlying population allele frequency is independent across populations and SNPs.

## Discussion

When looking at patterns of polygenic score overdispersion across populations, we observe highly inconsistent signals depending on the GWAS cohort from which we obtained effect size estimates. Because we are using the exact same population panels to obtain population allele frequencies in all tests, the source of the inconsistencies must necessarily come from differences in the effect size estimates in the different GWAS, or be due to different SNPs passing the significance threshold in different GWAS. These inconsistencies are not limited to tests involving height-associated SNPs: they also appear in tests involving SNPs associated with other phenotypes, like white blood cell count, mean corpuscular hemoglobin, potassium levels in urine, and educational attainment.

For those phenotypes for which we have effect size estimates from more than two different sources, we find that the GWASs performed using multiple cohorts of diverse ancestries - GIANT and PAGE - show stronger overdispersion in genetic scores via the *Q*_*X*_ statistic and stronger evidence of population stratification (Figures 4 and 5). In biobank-based GWAS conducted using panels with relatively homogeneous ancestries, the signals of selection are generally (but not always) more attenuated, and signals of stratification are much weaker. This suggests differences in scores are perhaps not driven by a biological signal and are instead driven by population stratification in GIANT and/or PAGE. An alternative explanation is that the overdispersion in PAGE or GIANT-derived scores is truly biological, and perhaps the GWAS performed in more homogeneous biobank studies are overcorrecting for population stratification, increasing the false negative rate of the *Q*_*X*_ statistic. Moreover, our power to find signals of polygenic adaptation might stem from SNPs with large contributions to phenotypic variance in non-European populations (e.g. African populations) and thus we might only be able to see these signals when we include individuals of African ancestry in a GWAS, as is done in PAGE.

Though plausible, alternative explanations seem less likely than stratification, but we cannot discard them at the moment, at least until a more extensive simulation study can allow us to compare these scenarios and their resulting score dispersion patterns. Another possible cause for these inconsistencies could be differences in the number of SNPs or individuals included in each GWAS, leading to differences in power to detect score overdispersion on trait-associated variants. Indeed, in some of the smaller cohorts (FINRISK and APCDR) we observe little to no evidence for strong deviations from neutrality in the distribution of genetic scores across populations.

We find that the type of test performed to obtain *Q*_*X*_ P-values does not yield strong differences in such P-values, at least not of the magnitude observed when using effect size estimates from different GWAS cohorts. Those phenotypes and GWAS cohorts for which we find significant overdispersion via the chi-squared distributional assumption for the *Q*_*X*_ statistic also tend to be the ones for which we find significant overdispersion when not relying on it. This suggests that this assumption - while not entirely accurate (Berg and Coop, 2014) - is still reasonably valid, across all the phenotypes we looked at, assuming the effect size estimates are not affected by stratification.

We were able to replicate the finding by Berg, Harpak, et al. (2019): there is a significant relationship between the differences in allele frequencies between GBR and other worldwide populations and the differences in effect size estimates between UKBB and GIANT. We note that a similar relationship was found by Uricchio et al. (2019), who showed an increase in the magnitude of allele frequency differences between GBR and TSI when ordering SNPs by their P-value in GIANT - an increase not observed when ordering them by their P-value in UKBB. We note, however, that this relationship is relatively absent in comparisons of UKBB and other GWAS, again suggesting that population stratification in GIANT, rather than over-correction of effect size estimates in UKBB, may be the culprit. In any case, Haworth et al. (2019), Novembre and NH Barton (2018), Coop (2019) and Rosenberg et al. (2019) encourage caution about the interpretation of signals of polygenic adaptation due to the presence of residual stratification even in GWAS panels with no clear evidence for stratification, as these signals may be subtle enough to escape notice, yet still affect this type of tests.

Furthermore, when we performed an artificial meta-analysis on the UKBB data, emulating the methodology of GIANT, we observed more dispersion of polygenic scores among populations than when using a single GWAS cohort, echoing findings by Kerminen et al. (2019) at a more localized geographic scale. As we previously observed in the vanilla (single-cohort) UKBB analysis, the less homogeneous the ancestries of the individuals in the cohort (“all ethnicities” vs. “white British”), the more dispersion is observed, which in turn causes a more inflated *Q*_*X*_ statistic. Nevertheless, both meta-analyses (“all ethnicities” and “white British”) show higher *Q*_*X*_ statistics than their single-cohort counterparts, regardless of the meta-analysis method deployed. This is also observed regardless of whether one uses cohort-specific PCs to correct for stratification in the meta-analysis or global PCs from a PCA including all individuals. Overall, this adds weight to the hypothesis that a failure of GWAS meta-analyses to control for population stratification may affect polygenic score tests against a neutral null hypothesis. We note that several of the component GWAS amalgamated in GIANT were not corrected via PCA or other standard methods of correction in common use today (Wood et al., 2014), so it is likely that we are being over-conservative in our simulations.

It is important to keep in mind that each particular GWAS used imposes strong conditions on the set of SNPs that are included in the *Q*_*X*_ analysis. We expect SNPs associated with a phenotype in a given cohort to explain more variance in the population from which that cohort was obtained than in other populations, simply because the significant SNPs need to have high enough allele frequencies in the study cohort for them to be recovered in the first place. It is unclear how this will affect false positive and false negative rates of tests of score overdispersion performed in different cohorts. For example, if we see a score overdispersion signal when using a GWAS from cohort 1 but not when using a GWAS from cohort 2, this could be due to a true positive and a lack of power in cohort 2, or due to an artefact caused by cohort 1. It is also possible that, if negative selection acts on some of the trait-related variation, it might affect statistical power by constraining large-effect alleles to be kept at lower frequencies, thus making large-effect alleles more population-specific.

Ultimately, the set of SNPs used in each analysis and their influence on the score depends on a complex combination of factors including allele frequencies, linkage disequilibrium with causal variants, statistical power for detection and effect size inflation due to the winner’s curse, together with the underlying evolutionary genetic process that “generates” the observed data. While modeling the individual effect of each of these factors on the inflation of the *Q*_*X*_ statistic is beyond the scope of this study, we note that all of these factors may be influencing the differences we observe among score sets. Indeed, when controlling for the set of SNPs included in the score, we see an attenuation of differences between scores (Figures S8 and S9).

In future studies of polygenic adaptation, we recommend the use of large homogeneous data sets and the verification of signals of polygenic score overdispersion in multiple GWAS cohorts (e.g. M Chen et al. (2019)). We also recommend caution even when finding that statistics testing against neutrality are significant in multiple GWAS cohorts: it is still possible that all the GWAS cohorts may be affected by subtle stratification or other confounding issues, possibly affecting different axes of population structure in different ways. To try to avoid stratification issues, recent studies have proposed to look for evidence for polygenic adaptation within the same panel that was used to obtain SNP effect size estimates, i.e. avoiding comparisons between populations that might be made up of individuals outside of the GWAS used to obtain effect size estimates (e.g. X Liu et al. (2018)). The argument favored by these studies is that, by ensuring that the population on which the GWAS was performed and from which allele frequencies are obtained matches exactly, one need not be confounded by differences in estimates between these populations (for example, due to gene-by-environment interactions). However, Mostafavi et al. (2019) recently showed that the accuracy of polygenic scores often depends on the age and sex composition of the GWAS study participants, even when studying individuals of roughly similar ancestries within a single cohort, due to heritability differences along these axes of variation. This implies that ancestry-based stratification is not the only confounder that researchers should be aware of when trying to detect polygenic adaptation.

Approaches based on tree sequence reconstructions along the genome (Hubisz and Siepel, 2020; Kelleher et al., 2016; Rasmussen et al., 2014; Speidel et al., 2019) appear to be a fruitful avenue of research towards the development of methods that can properly control for some of these confounders. These methods can model local genealogical relationships among individuals, which can in turn serve to track the segregation of trait-associated alleles backwards in time. For example, Stern et al. (2020) recently showed that a method for detecting polygenic adaptation based on tree sequences is highly robust to GWAS stratification, ascertainment bias in SNP effects and negative selection, among other potential confounders. They were also able to show that the signal of polygenic adaptation previously found at educational attainment-associated variants may be due to indirect selection on other, correlated, traits.

Overall, we generally urge caution in the interpretation of signals of polygenic score overdispersion based on human GWAS data, at least until we have robust generative models that can explain how stratification is creeping into these tests (Young et al., 2019). This is especially important when working with socially-charged traits like educational attainment, which are rife for misuse and misinterpretation, and potentially affected by unaccounted socioeconomic and cultural confounding factors. Due to the high risk of misappropriation of this type of results by hate groups (Harmon, 2018), we also recommend that researchers make an effort to explain the caveats and problems associated with these tests in their publications (Coop, 2019; Novembre and NH Barton, 2018; Rosenberg et al., 2019), as well as the strong sensitivity of their performance to the input datasets that we choose to feed into them.

## Supplementary material

The code used to perform the analyses in this manuscript is available at: https://github.com/albarema/GWAS_choice/.

## Acknowledgments

We thank Jeremy Berg, Lawrence Uricchio, Mashaal Sohail, Bárbara Bitarello, Sriram Sankararaman and Arun Durvasula for helpful comments on the manuscript, as well as Mark Daly, Samuli Ripatti, Jukka Koskela, Masahiro Kanai, Yukinori Okada, Yoichiro Kamatani, Evan Irving-Pease and Graham Gower for general advice and technical assistance in obtaining and handling association data. We would also like to thank the members of the Racimo group for feedback throughout the duration of the project. Finally, we thank all participants of the biobanks and association studies included in this work, for their valuable contribution to the study of genetic variation and disease. Access to the UK Biobank data was provided to AA via application number 32683 and to ARM via application number 31063. FR is supported by a Villum Fonden Young Investigator award (project no. 00025300). AR-M is supported by the Lundbeck Foundation GeoGenetics Centre grant (R302-2018-2155) and a Novo Nordisk grant to the GeoGenetics Centre (NNF18SA0035006). ARM is supported by funding from the National Institutes of Health (K99MH117229). Version 5 of this preprint has been peer-reviewed and recommended by Peer Community In Evolutionary Biology (https://doi.org/10.24072/pci.evolbiol.100125).

## Conflicts of interest disclosure

The authors of this preprint declare that they have no financial conflict of interest with the content of this article. Fernando Racimo is a PCI EvolBiol recommender.

## Supplementary Figures

**Figure S1.**
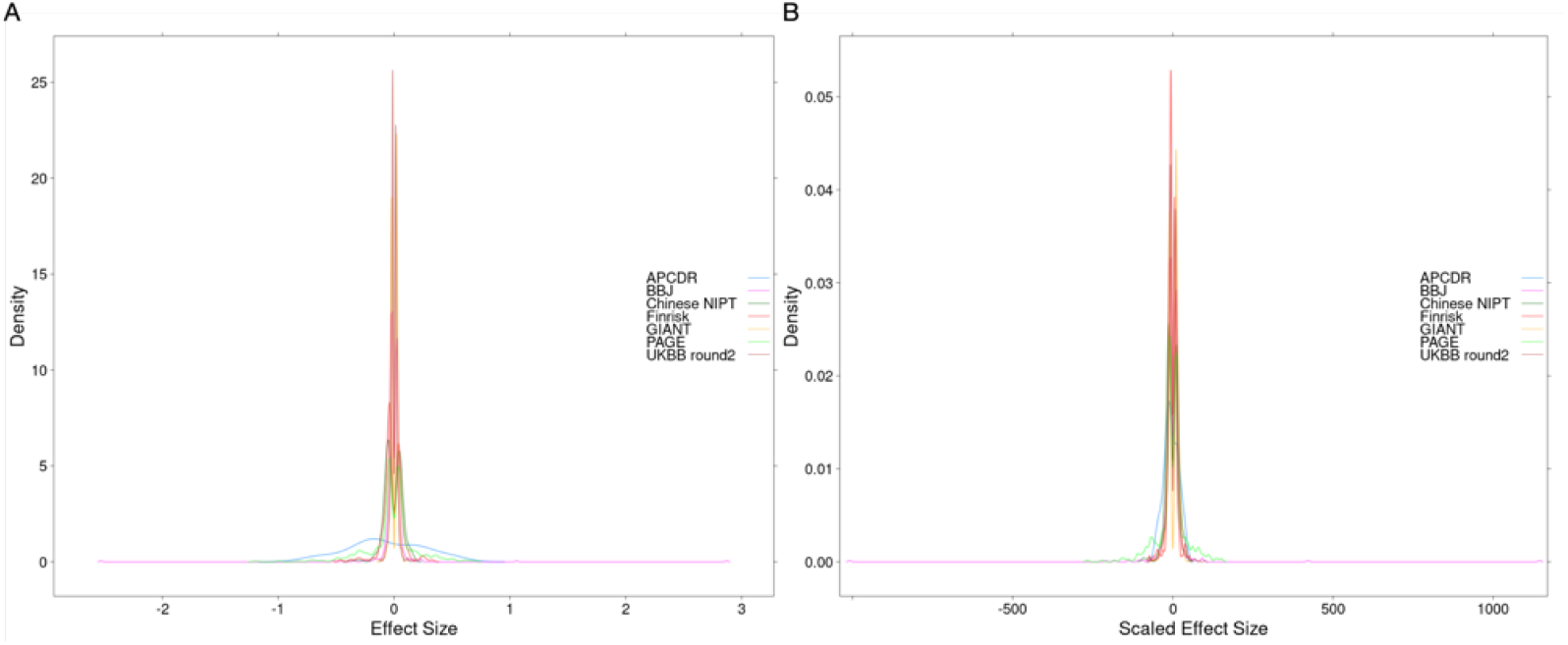
A. Distribution of effect size estimates (with respect to the derived allele) for approximately independent height-associated SNPs with *P* <1_e_−5. B. Distribution of the product of the effect size estimates and the square root of the sample size of the study from which they were obtained, for the same set of SNPs as in panel A.

**Figure S2.**
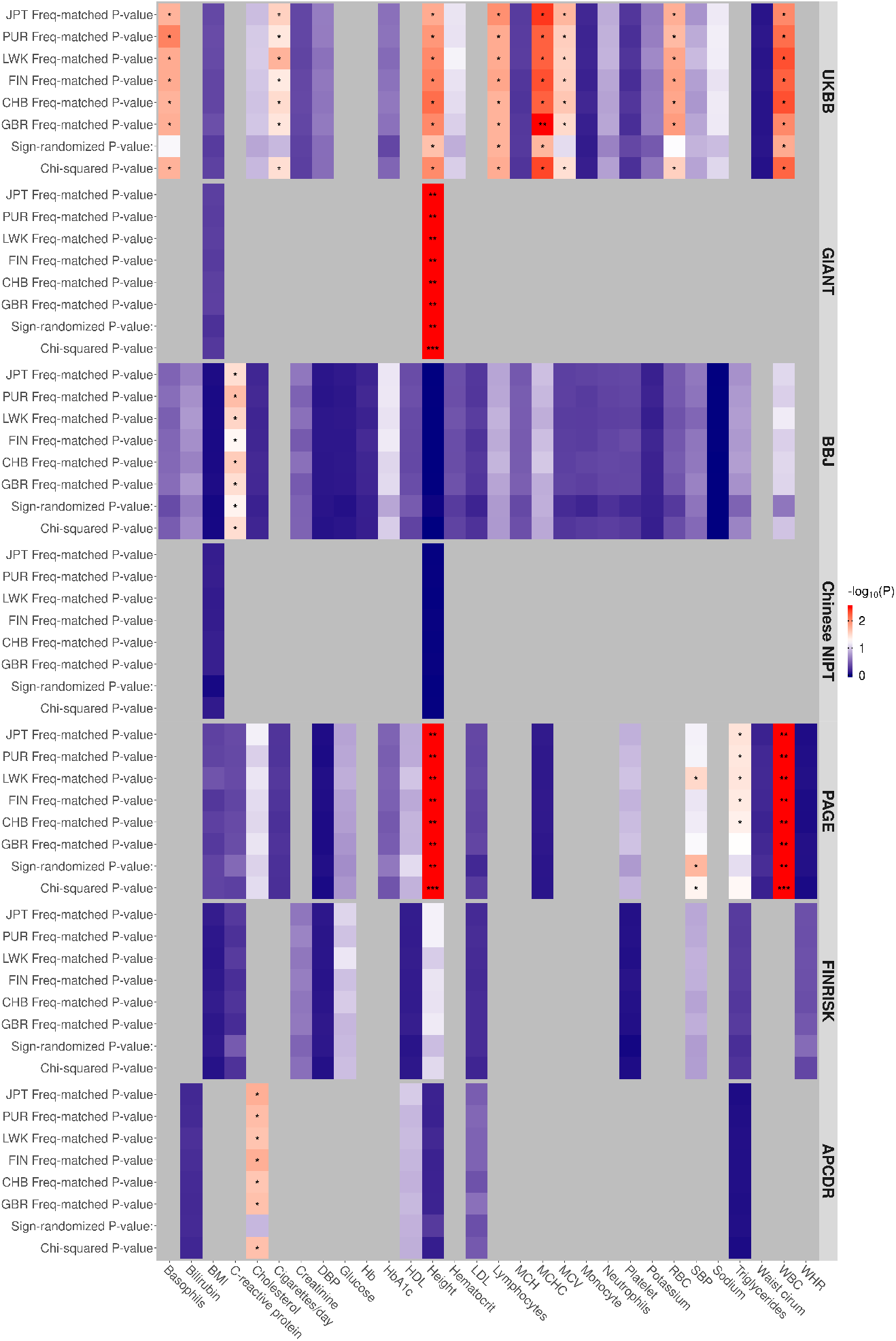
Evidence for polygenic score overdispersion on trait-associated SNPs (−*log*_10_(P-value)), using the *Q*_*X*_ test statistic. Each row in the heatmap corresponds to a specific GWAS cohort and a specific type of scheme to determine the significance of the *Q*_*X*_ statistic. The columns correspond to the different traits for each GWAS cohort that have SNPs with a P-value lower than 1_e_−5. “Freq-matched P-value” = P-value obtained by sampling SNPs with matching frequencies to the trait-associated SNPs in a particular cohort. “Sign-randomized P-value” = P-value obtained by randomizing the signs of the effect size estimates. “Chi-squared P-value” = P-value obtained by assuming the *Q*_*X*_ statistics has a chi-squared distribution.

**Figure S3.**
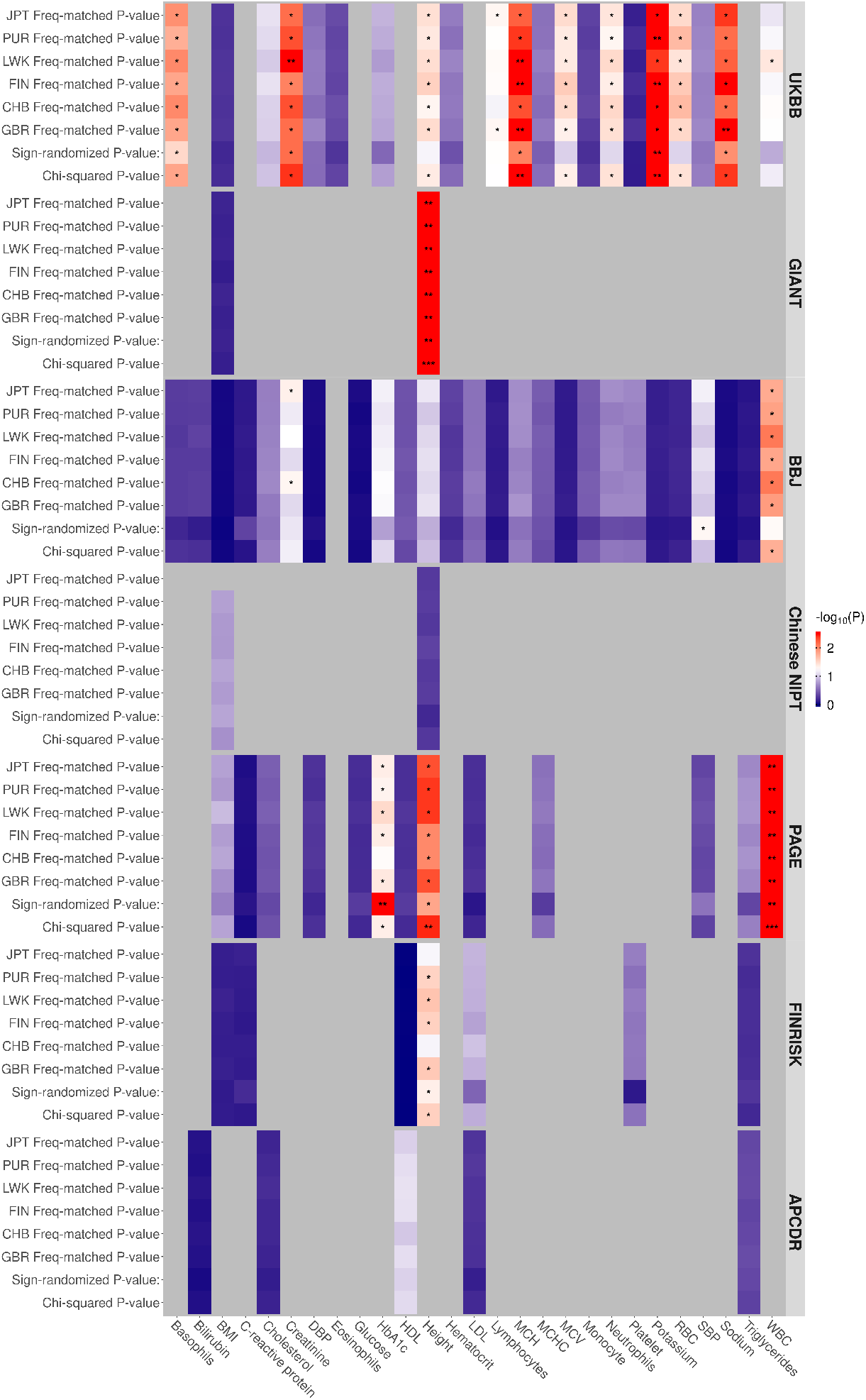
Evidence for polygenic score overdispersion on trait-associated SNPs (−*log*_10_(P-value)), using the *Q*_*X*_ test statistic. Each row in the heatmap corresponds to a specific GWAS cohort and a specific type of scheme to determine the significance of the *Q*_*X*_ statistic. The columns correspond to the different traits for each GWAS cohort that have SNPs with a P-value lower than 5_e_−8. “Freq-matched P-value” = P-value obtained by sampling SNPs with matching frequencies to the trait-associated SNPs in a particular cohort. “Sign-randomized P-value” = P-value obtained by randomizing the signs of the effect size estimates. “Chi-squared P-value” = P-value obtained by assuming the *Q*_*X*_ statistics has a chi-squared distribution.

**Figure S4.**
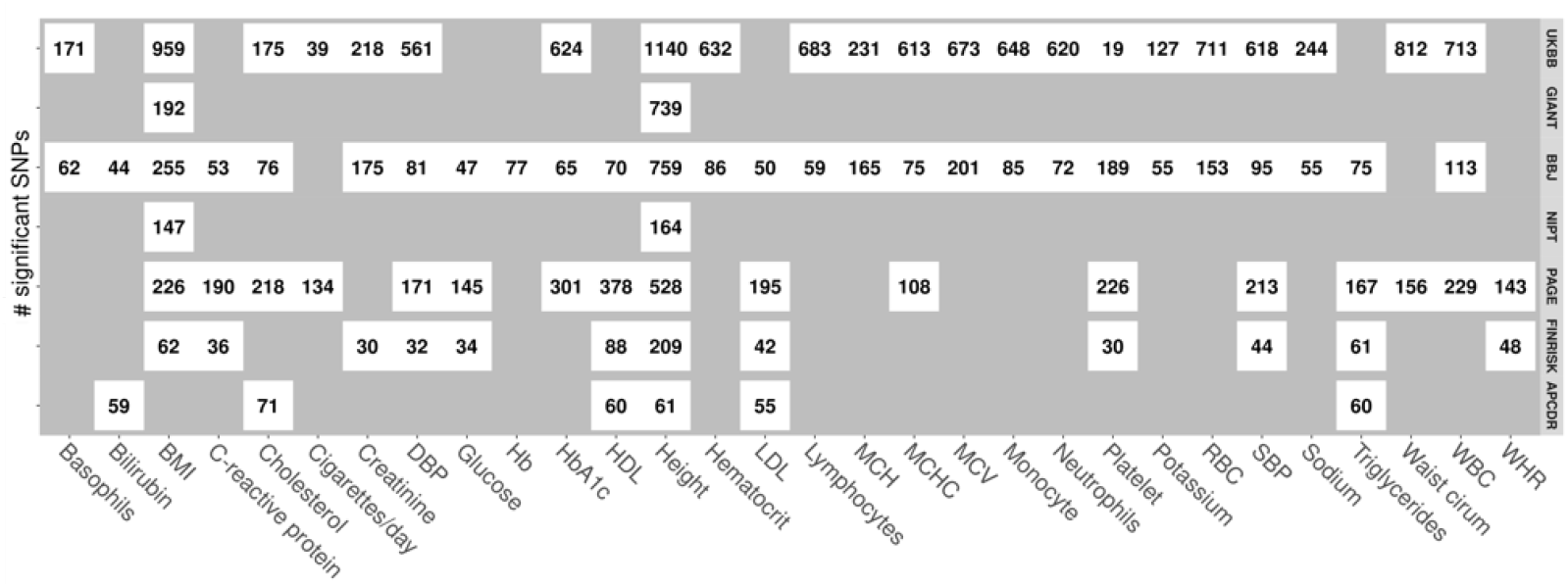
Number of significant SNPs used to compute the *Q*_*X*_ statistics for each cell of Figure 2. *P* < 1_e_−5.

**Figure S5.**
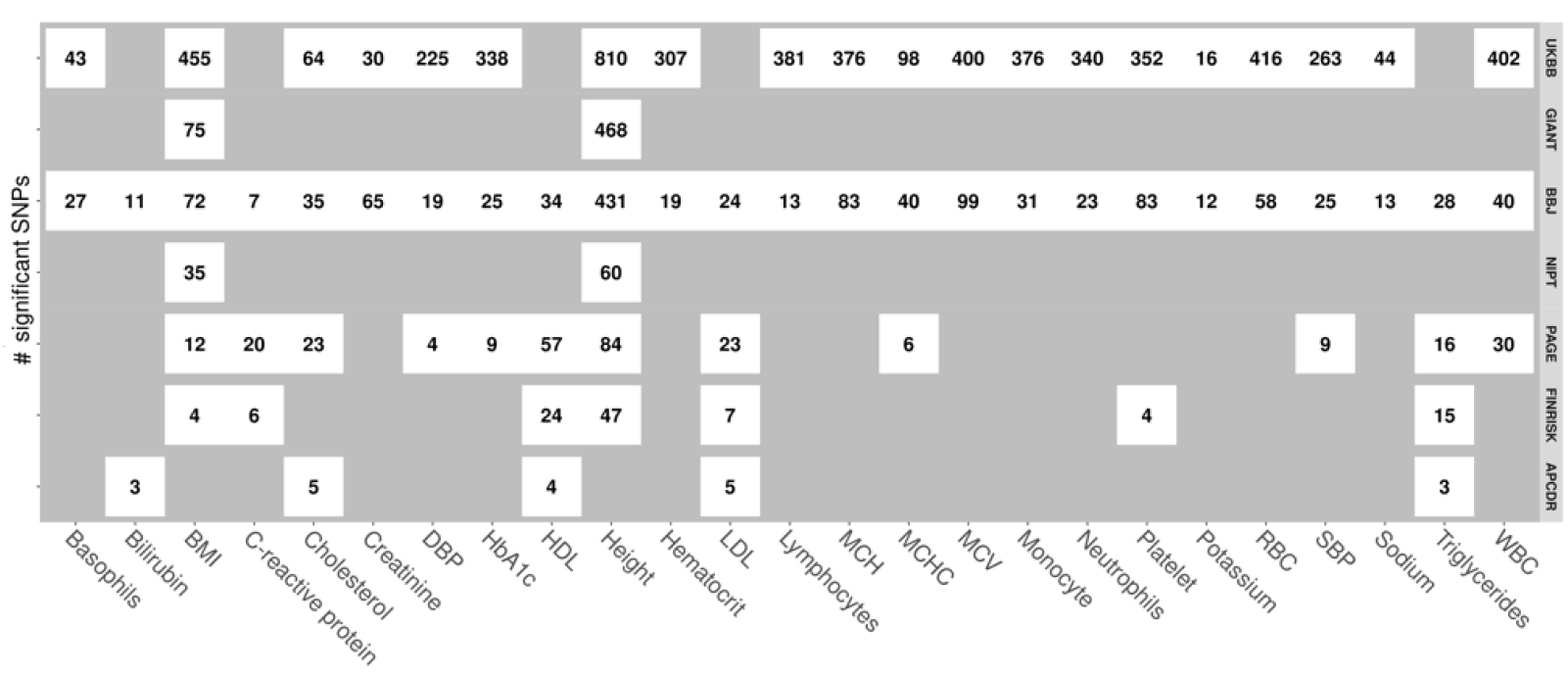
Number of significant SNPs for each trait in each GWAS with *P* < 5_e_−8.

**Figure S6.**
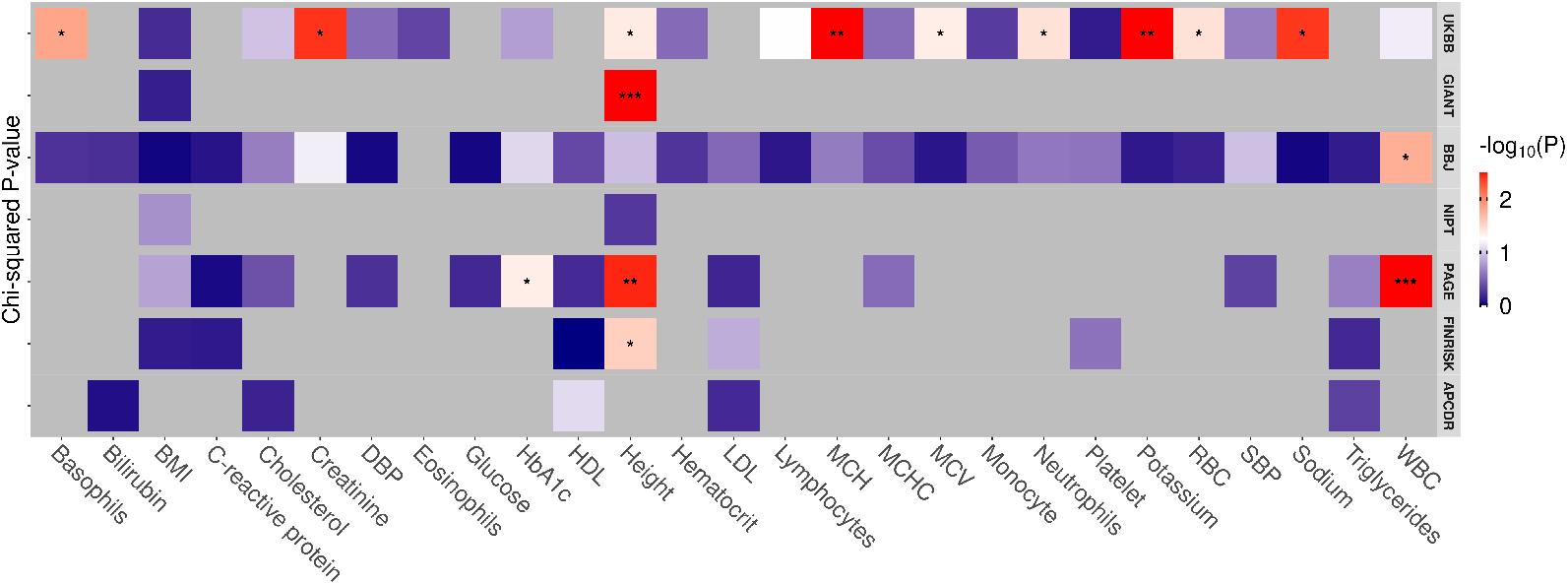
Evidence for polygenic score overdispersion on trait-associated SNPs (−*log*_10_(P-value)), using the *Q*_*X*_ test statistic. Each row in the heatmap corresponds to one of the six GWAS cohorts we are evaluating and the columns correspond to the different traits for each GWAS cohort that have SNPs with a P-value lower than 5_e_−8. BMI, body mass index; DBP, Diastolic blood pressure; HbA1c, glycated hemoglobin; HDL, high-density lipoprotein; LDL, low-density lipoprotein; MCH, mean corpuscular hemoglobin; MCHC, mean corpuscular hemoglobin concentration; MCV, mean corpuscular volume; RBC, red blood cell count; SBP, systolic blood pressure; WBC, white blood cell count; WHR, waist-to-hip ratio. Significance thresholds after Bonferroni corrections: *** denotes P < 0.05/m, ** denotes P < 0.05/n, * denotes P < 0.05 where *n* is the number of traits measured in each GWAS (row-dependent) and *m* is the total number of tests calculated, across all GWAS.

**Figure S7.**
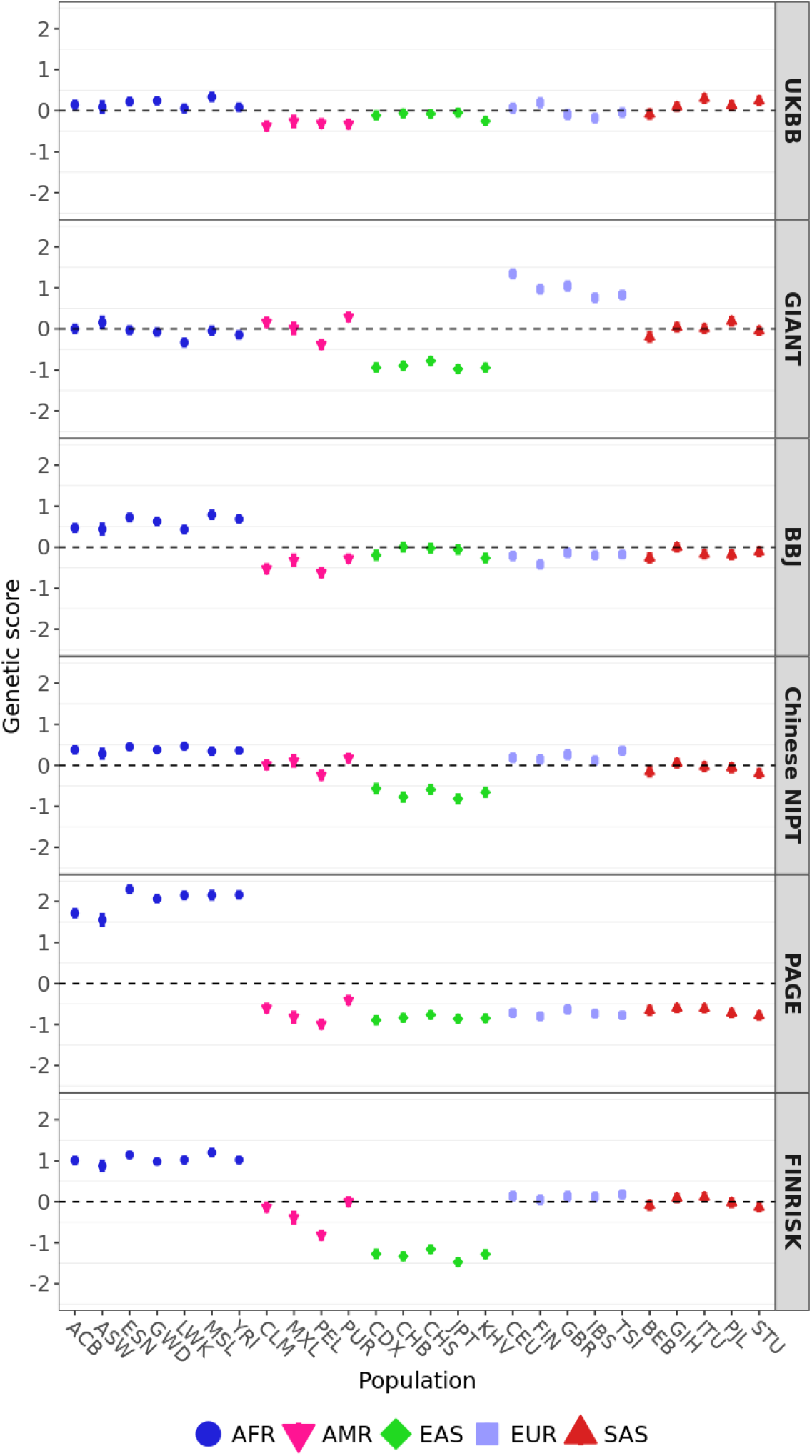
Polygenic scores for height using candidate SNPs with *P* < 5_e_−8 in 1000 Genome populations colored by their super-population code. The corresponding number of trait-associated SNPs and the *Q*_*X*_ P-value for each GWAS are shown in the bottom row of Table 3. Error bars denote 95% credible intervals, constructed using the method in Sohail et al., 2019, assuming that the posterior distribution of the underlying population allele frequency is independent across populations and SNPs.

**Figure S8.**
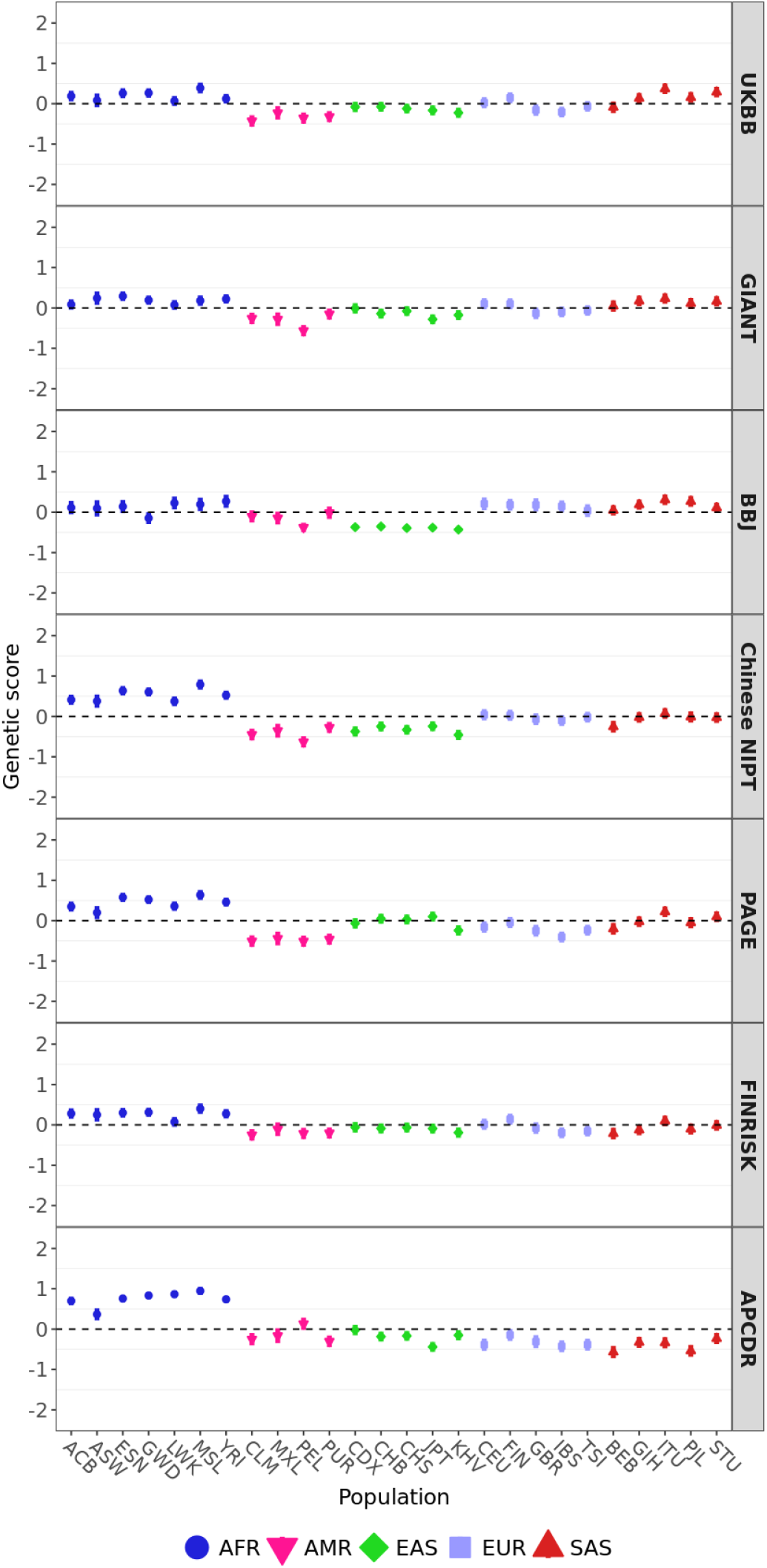
Polygenic scores for height 1000 Genome populations colored by their super-population code. Candidate SNPs were ascertained in the UKBB using the cutoff of *P* < 1_e_−5. Error bars denote 95% credible intervals, constructed using the method in Sohail et al., 2019, assuming that the posterior distribution of the underlying population allele frequency is independent across populations and SNPs.

**Figure S9.**
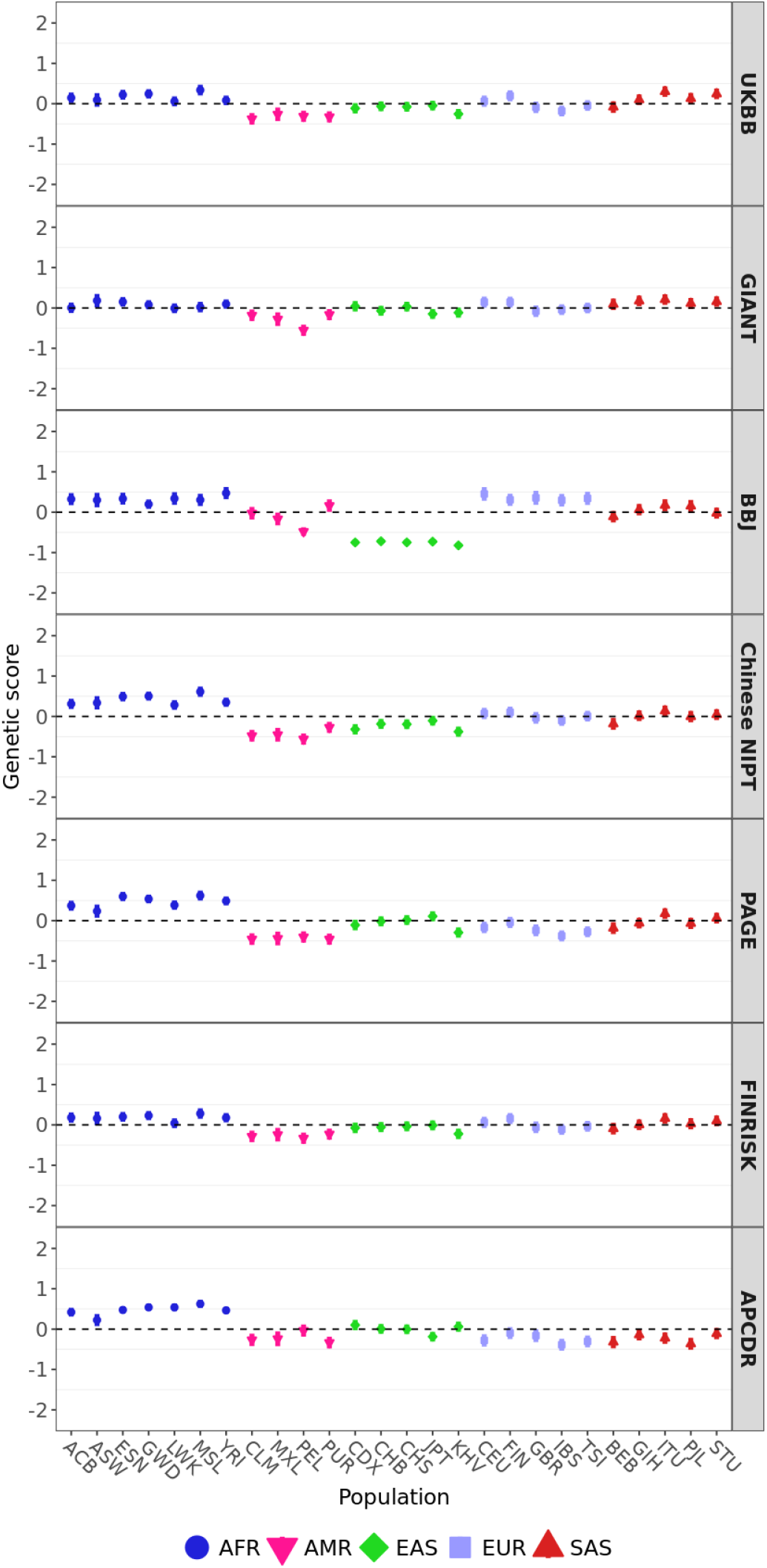
Polygenic scores for height 1000 Genome populations colored by their super-population code. Candidate SNPs were ascertained in the UKBB using the cutoff of *P* < 5_e_−8. Error bars denote 95% credible intervals, constructed using the method in Sohail et al., 2019, assuming that the posterior distribution of the underlying population allele frequency is independent across populations and SNPs.

**Figure S10.**
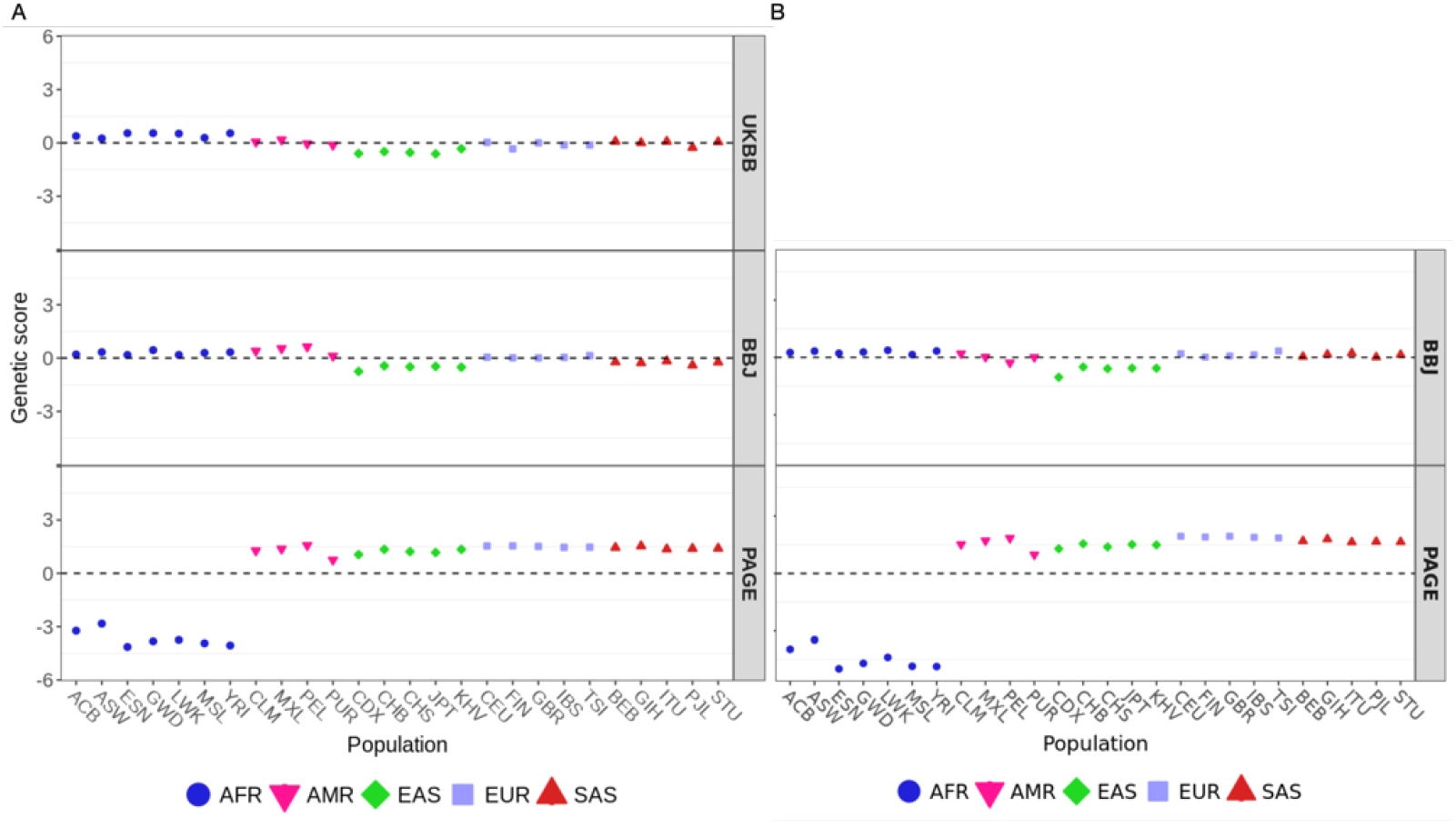
Polygenic scores for white blood cell counts using candidate SNPs with A. *P* < 1e−5 and B. *P* < 5_e_−8 in the 1000 Genomes Project populations colored by their super-population code. Error bars denote 95% credible intervals, constructed using the method in Sohail et al., 2019, assuming that the posterior distribution of the underlying population allele frequency is independent across populations and SNPs.

**Figure S11.**
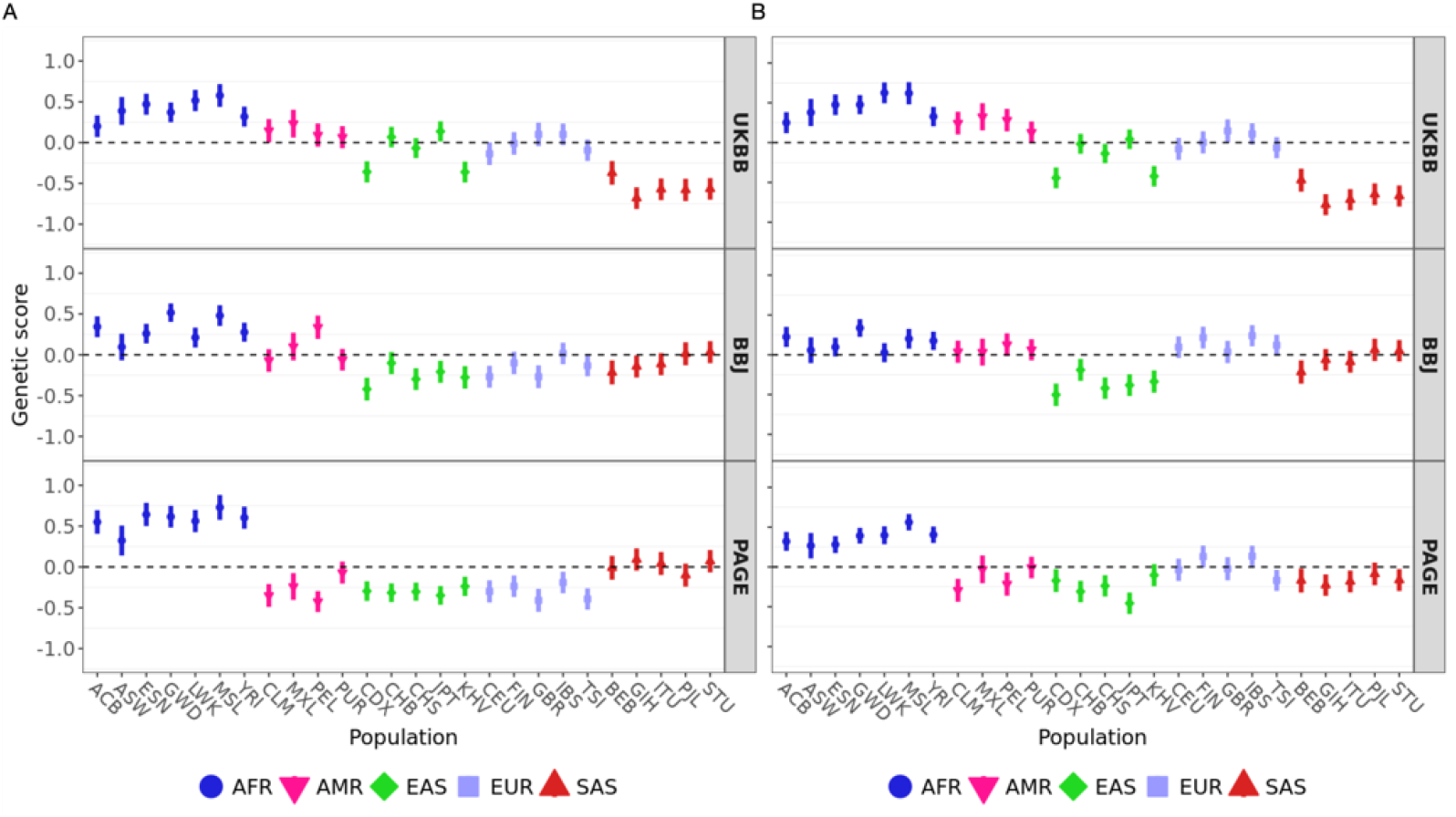
Polygenic scores for mean corpuscular hemoglobin using candidate SNPs with A. *P* < 1_e_−5 and B. *P* < 5_e_−8 in 1000 Genome populations colored by their super-population code. X trait-associated SNPs. Error bars denote 95% credible intervals, constructed using the method in Sohail et al., 2019, assuming that the posterior distribution of the underlying population allele frequency is independent across populations and SNPs.

**Figure S12.**
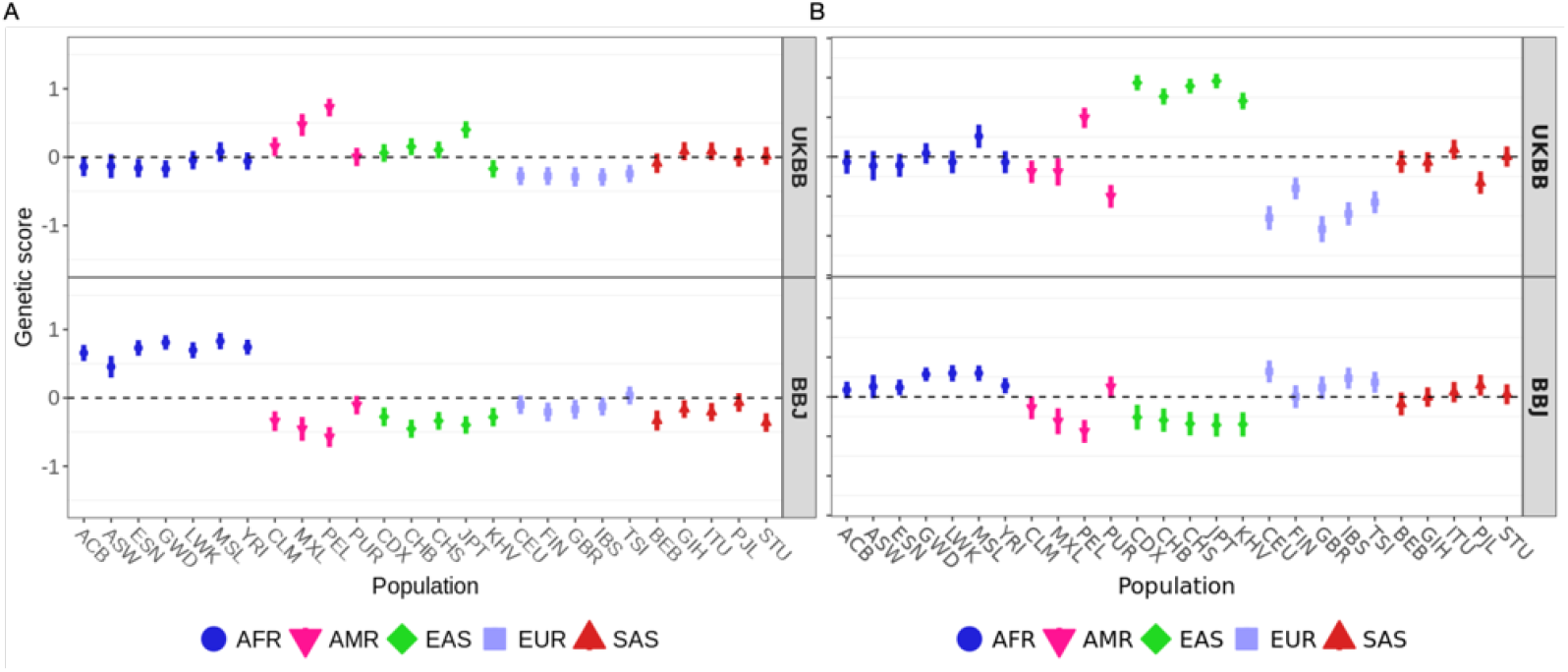
Polygenic scores for potassium level using candidate SNPs with A. *P* < 1_e_−5 and B. *P* < 5_e_−8 in 1000 Genome populations colored by their super-population code. Error bars denote 95% credible intervals, constructed using the method in Sohail et al., 2019, assuming that the posterior distribution of the underlying population allele frequency is independent across populations and SNPs.

**Figure S13.**
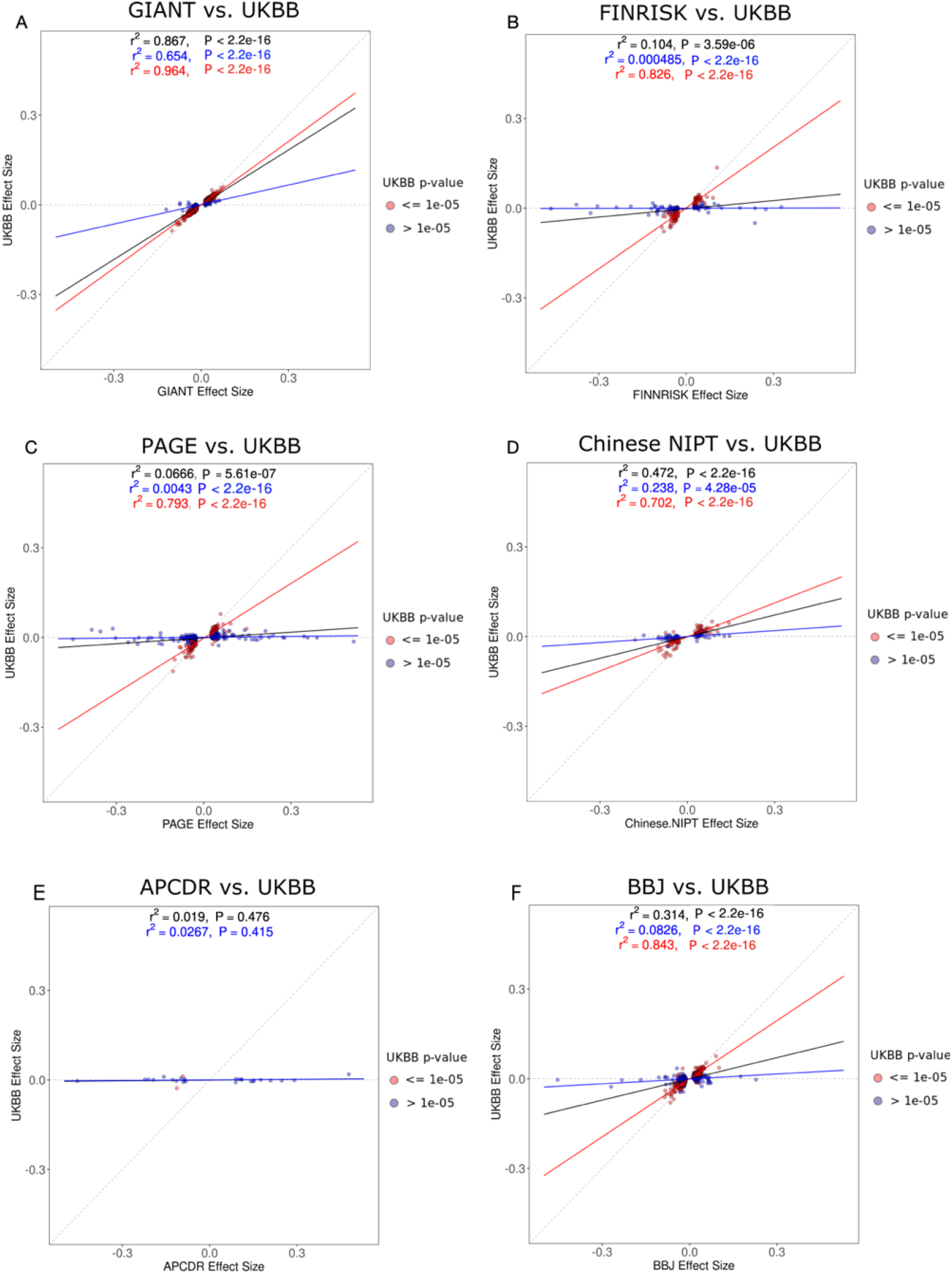
Regression of UKBB height effect size estimates against non-UKBB height effect size estimates, after filtering for SNPs with *P* < 1_e_−5 in the non-UKBB GWAS. The SNPs are colored based on their P-value in the UKBB, as are the corresponding regression lines (red: *P* < 1_e_−5; blue: *P* > 1_e_−5). The black regression line was obtained using all SNPs, regardless of their P-value in the UKBB GWAS.

**Figure S14.**
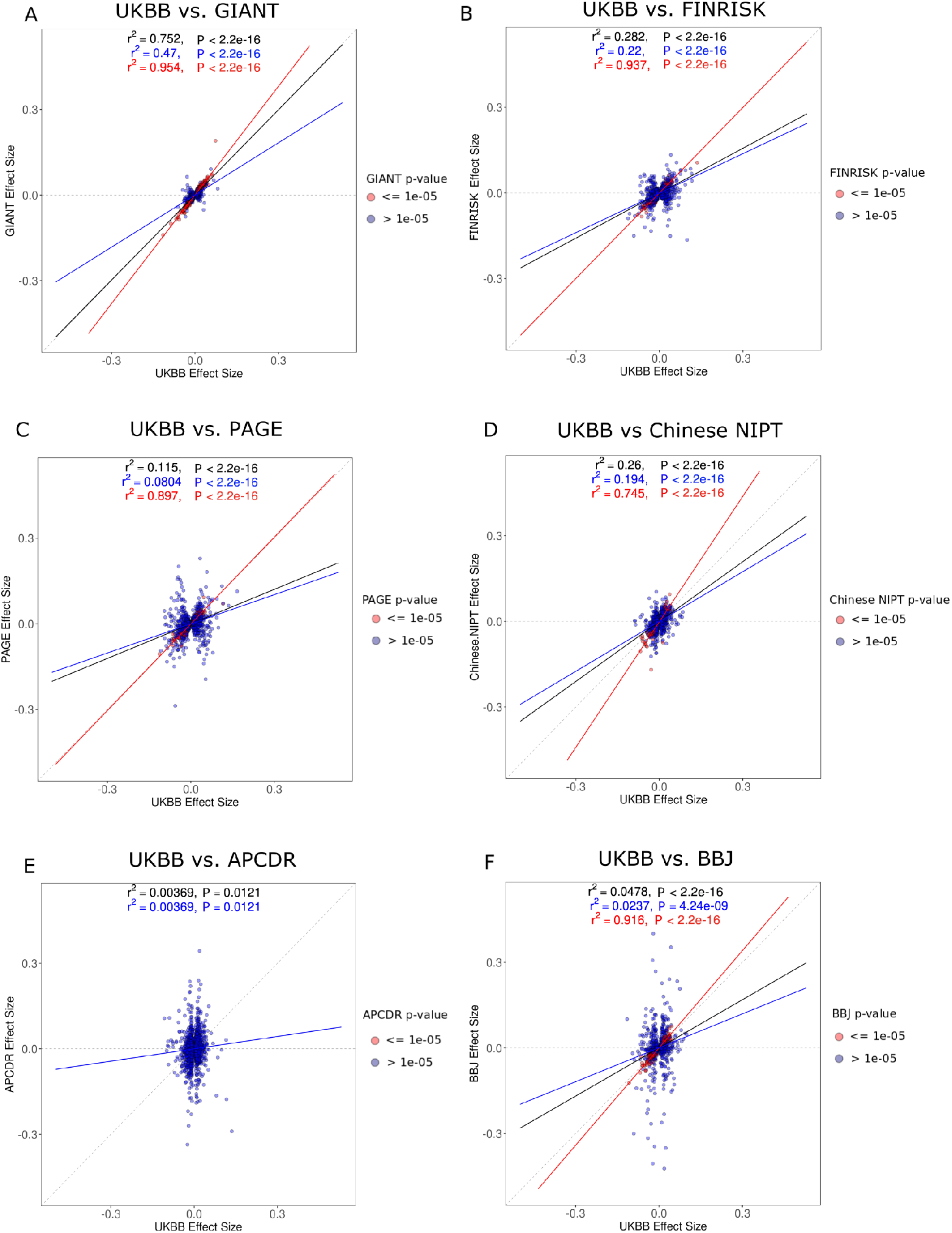
Regression of non-UKBB height effect size estimates against UKBB height effect size estimates, after filtering for the SNPs with the lowest UKBB P-values in 1,703 approximately-independent LD blocks. The SNPs are colored based on their P-value in the non-UKBB GWAS, as are the corresponding regression lines (red: *P* < 1_e_−5; blue: *P* > 1_e_−5). The black regression line was obtained using all SNPs, regardless of their P-value in the non-UKBB GWAS.

**Figure S15.**
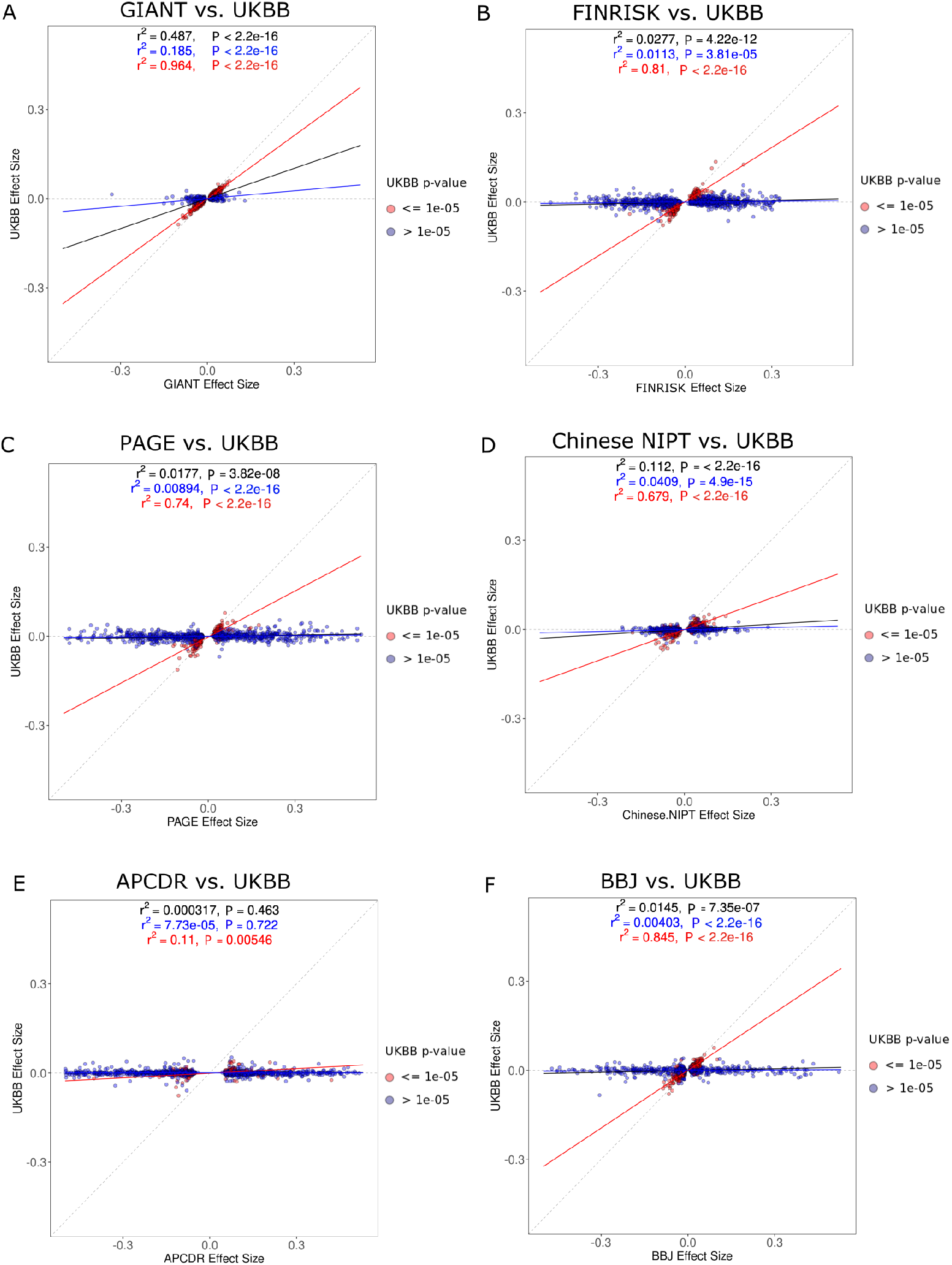
Regression of UKBB height effect size estimates against non-UKBB height effect size estimates, after filtering for the SNPs with the lowest non-UKBB P-values in 1,703 approximately-independent LD blocks. The SNPs are colored based on their P-value in the UKBB GWAS, as are the corresponding regression lines (red: *P* < 1_e_−5; blue: *P* > 1_e_−5). The black regression line was obtained using all SNPs, regardless of their P-value in the UKBB GWAS.

**Figure S16.**
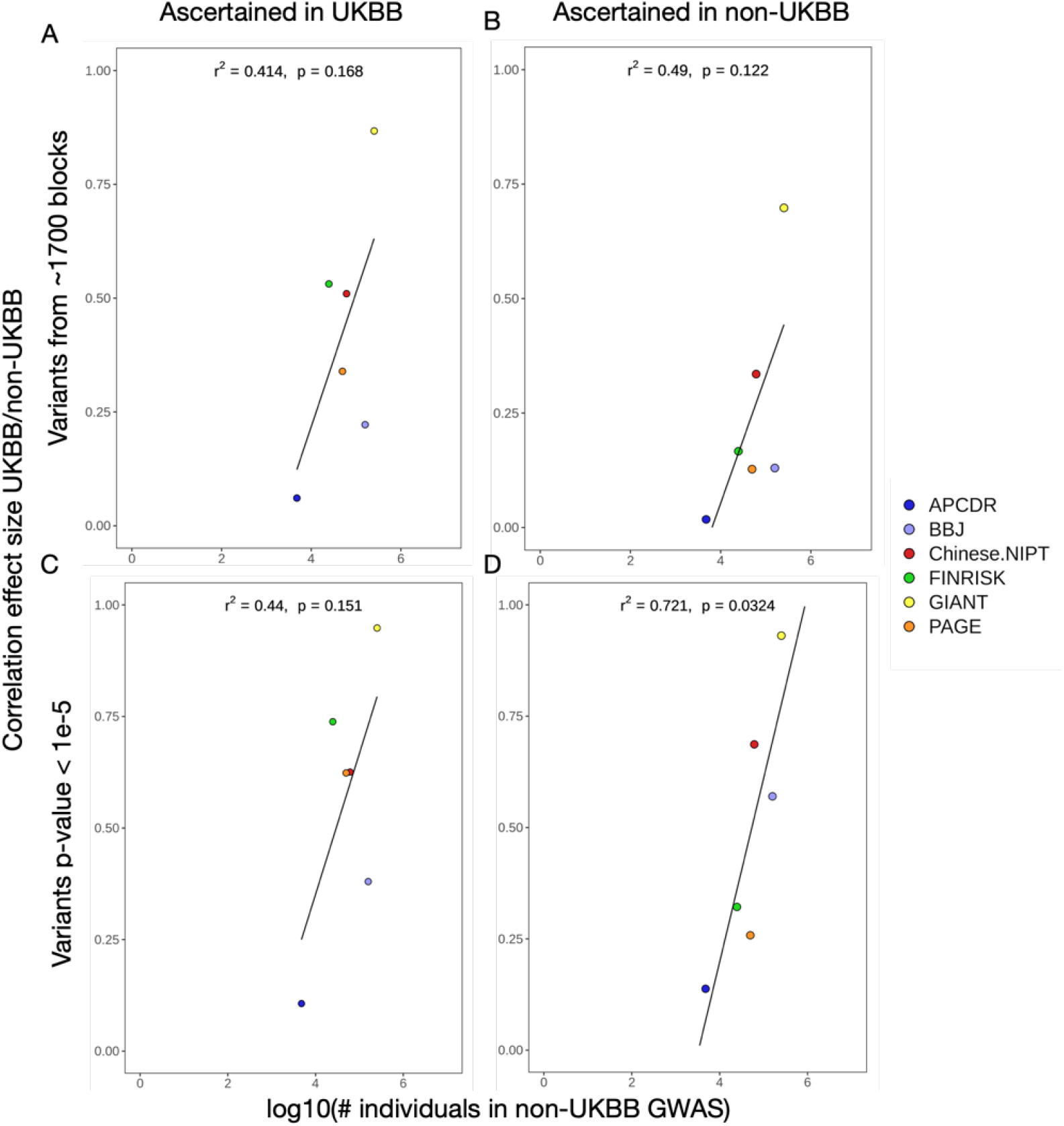
Correlation between the number of individuals in a GWAS and the correlation of effect size estimates between two different GWAS. In all panels, we compare UKBB against one of the other studies. **A and B**. Coefficients obtained when we include the SNPs from the 1,703 LD blocks. **C and D**. Coefficients obtained when we filter out those SNPs with a P-value above the threshold (*P* > 1_e_−5).

**Figure S17.**
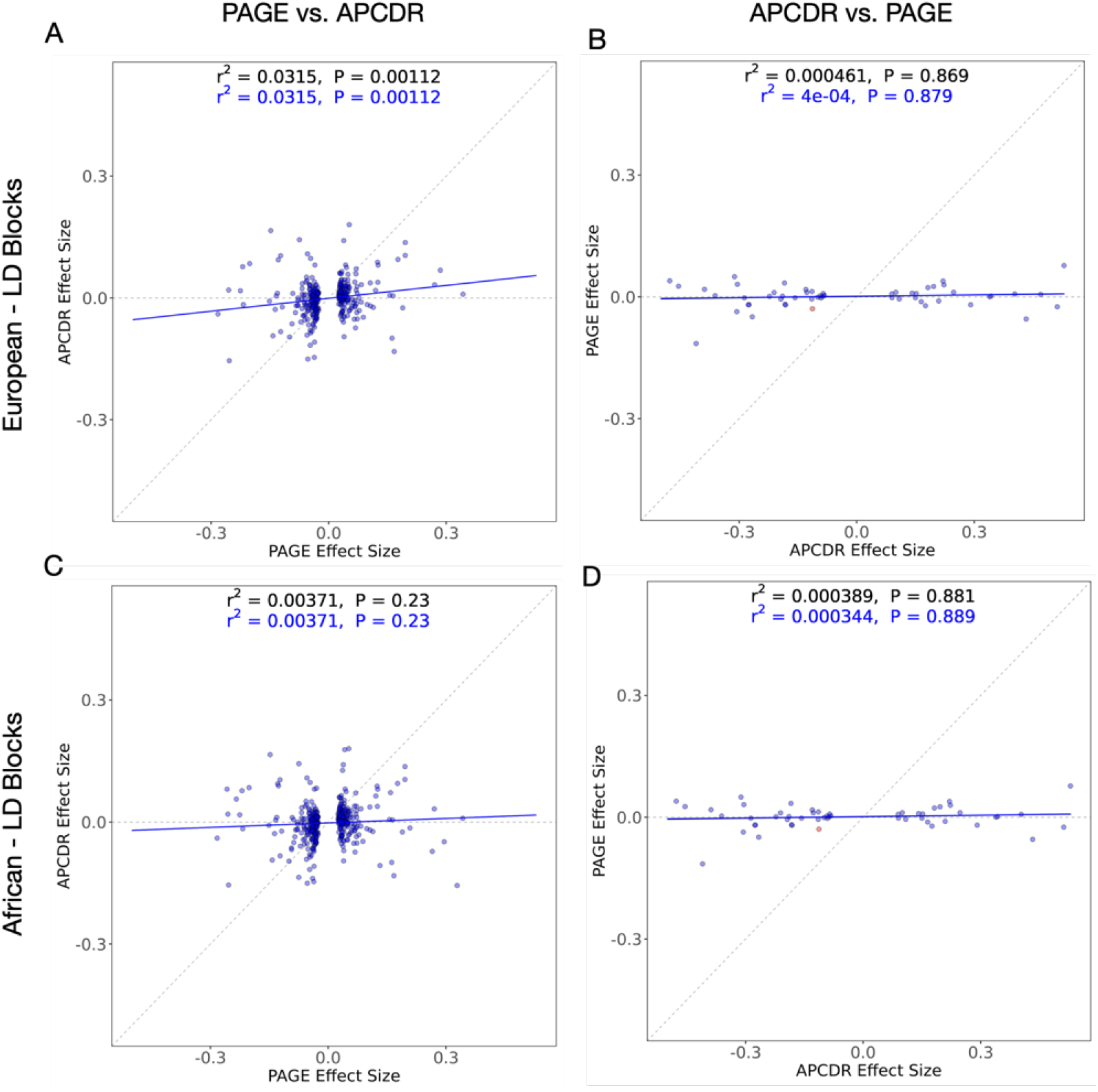
Regression of effect size estimates obtained from PAGE and APCDR. In the top panels (A and B), we used European LD blocks and in the lower panels (C and D) we used African LD blocks. In the left panel (A and C) we ascertained significant SNPs based on their P-values in the PAGE GWAS (*P* < 1_e_−5). In the right panel (B and D), we ascertained significant SNPs based on their P-values in the APCDR GWAS (*P* < 1_e_−5). The SNPs are colored based on their P-value in the non-ascertained GWAS, as are the corresponding regression lines (red: *P* < 1_e_−5; blue: *P* > 1_e_−5). The black regression line was obtained using all SNPs, regardless of their P-value in the non-ascertained GWAS.

**Figure S18.**
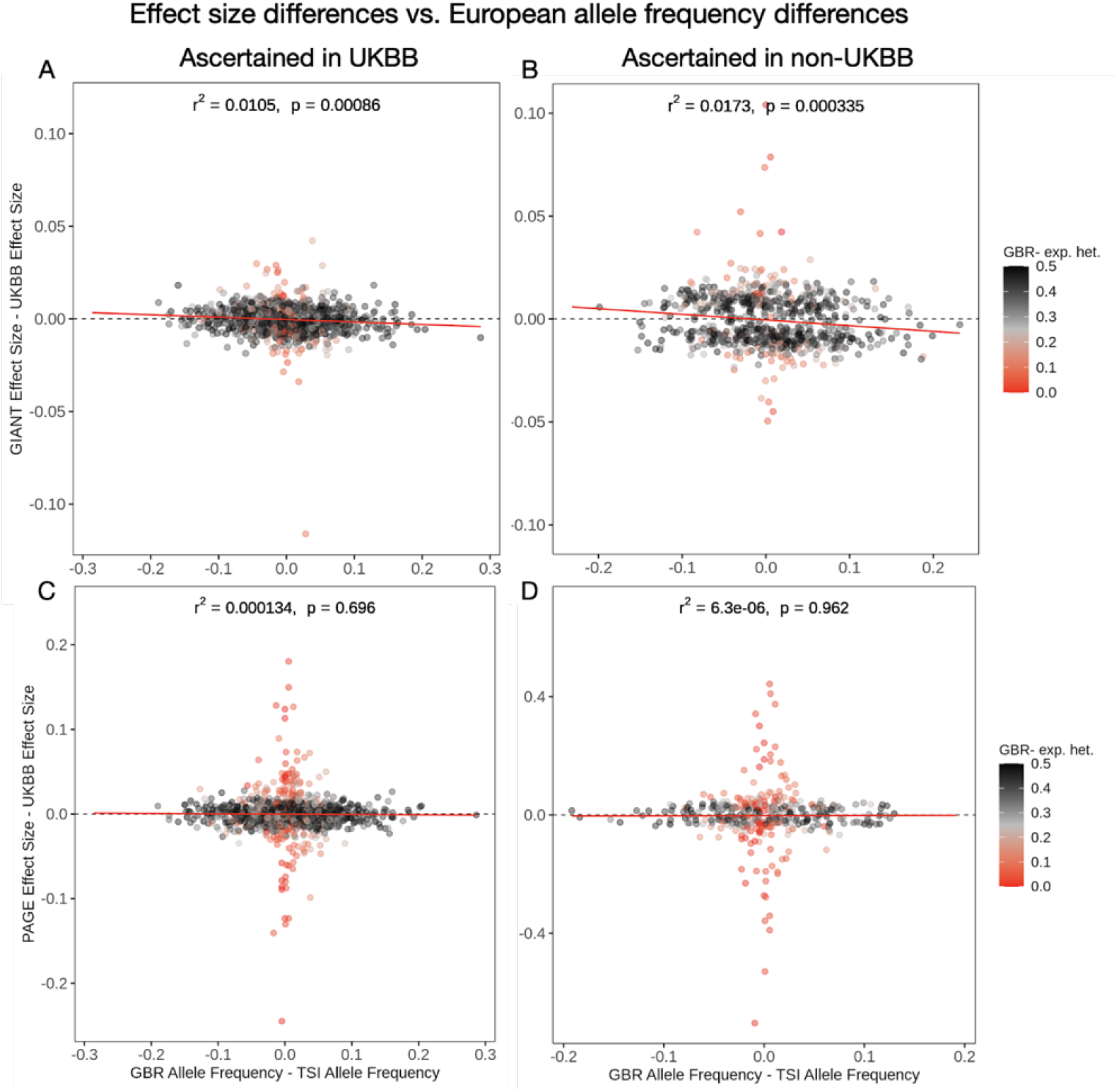
Regression of GWAS height effect size differences against allele frequency differences between northern and southern European population panels (GBR and TSI). We selected SNPs for this analysis that had P < 1_e_−5. SNPs are colored by their expected heterozygosity (2p(1-p)) in the GBR population.

**Figure S19.**
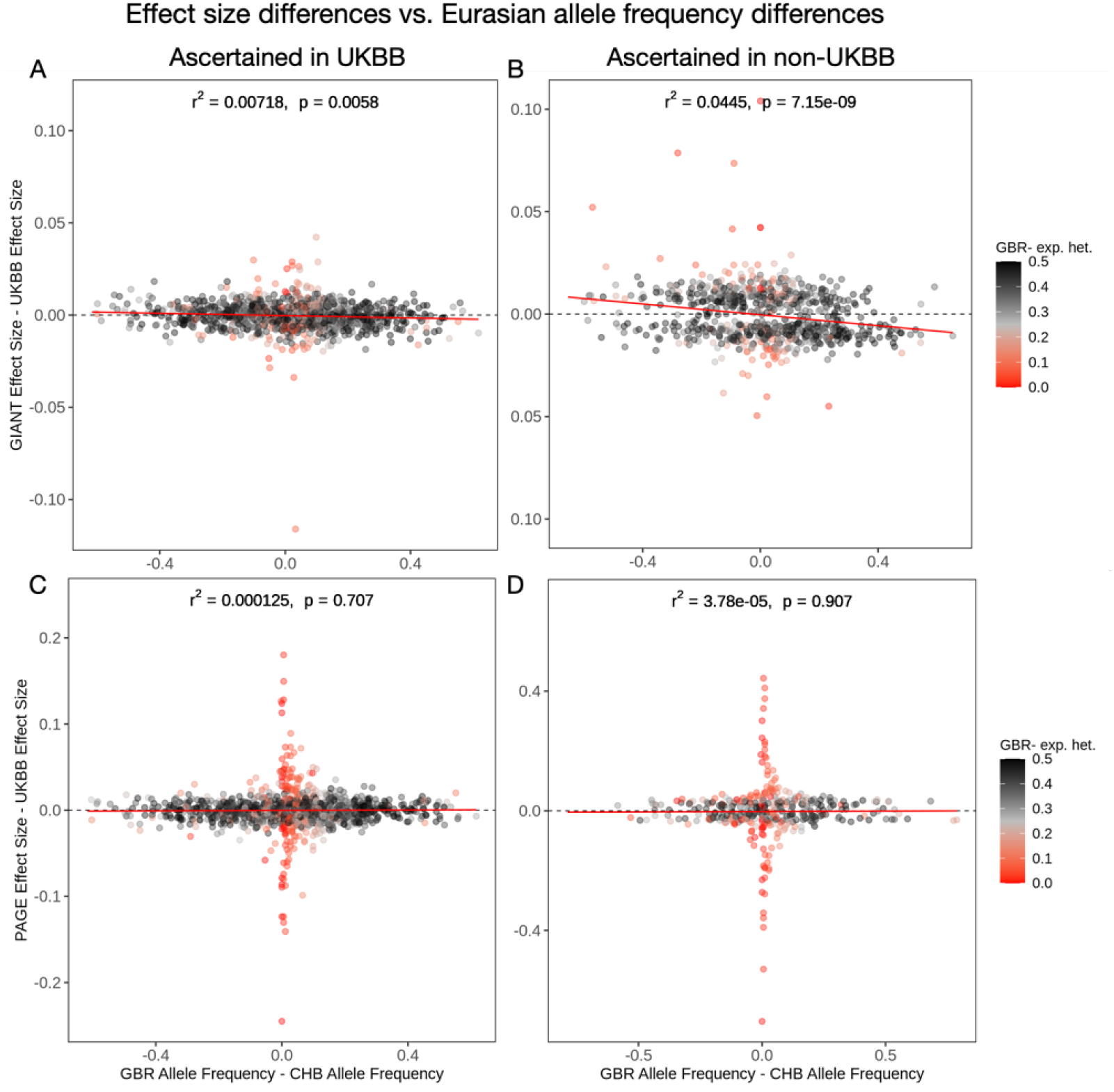
Each panel in this figure shows the same regressions as Figure S18, except that the difference in allele frequency is between a European and Asian population panels (GBR and CHB). SNPs are colored by their expected heterozygosity (2p(1-p)) in the GBR population.

**Figure S20.**
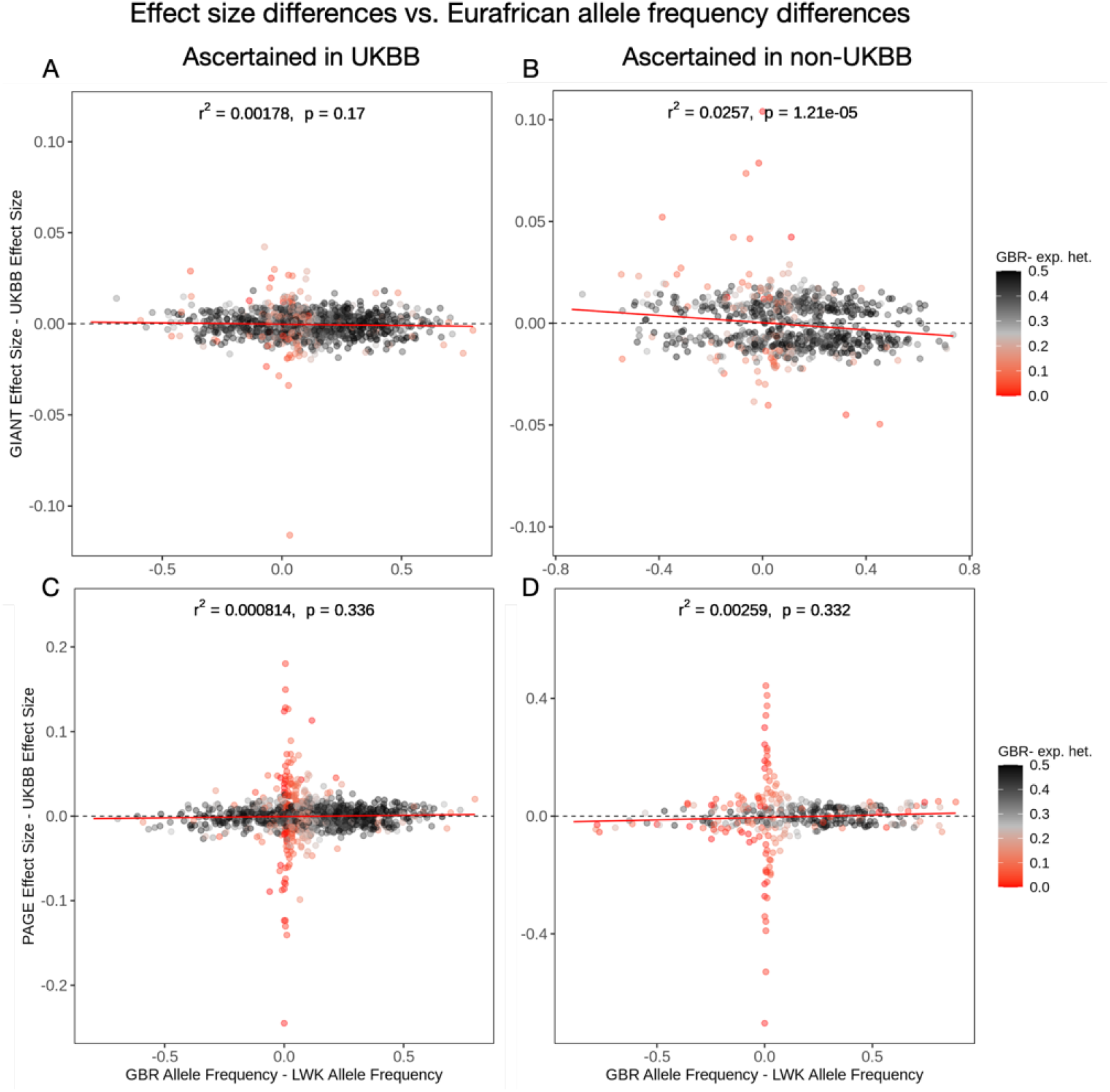
Each panel in this figure shows the same regressions as Figure S18, except that the difference in allele frequency is between the European and African populations panels (GBR and LWK).SNPs are colored by their expected heterozygosity (2p(1-p)) in the GBR population.

**Figure S21.**
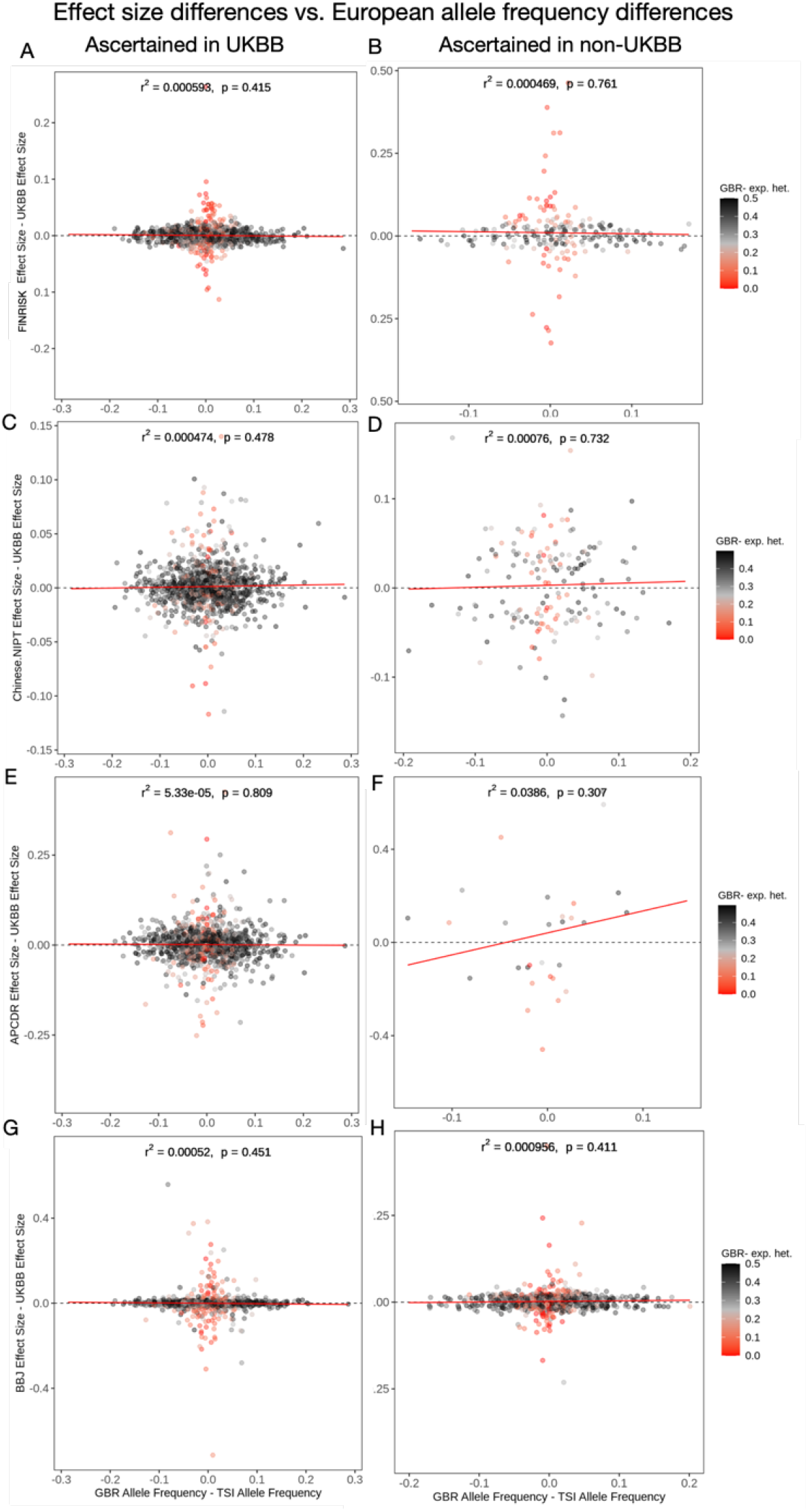
Regression of GWAS height effect size differences against allele frequency differences between the northern and southern European populations (GBR and TSI). **A**,**B**. We selected SNPs for this analysis that had P < 1_e_−5. SNPs are colored by their expected heterozygosity (2p(1-p)) in the GBR population.

**Figure S23.**
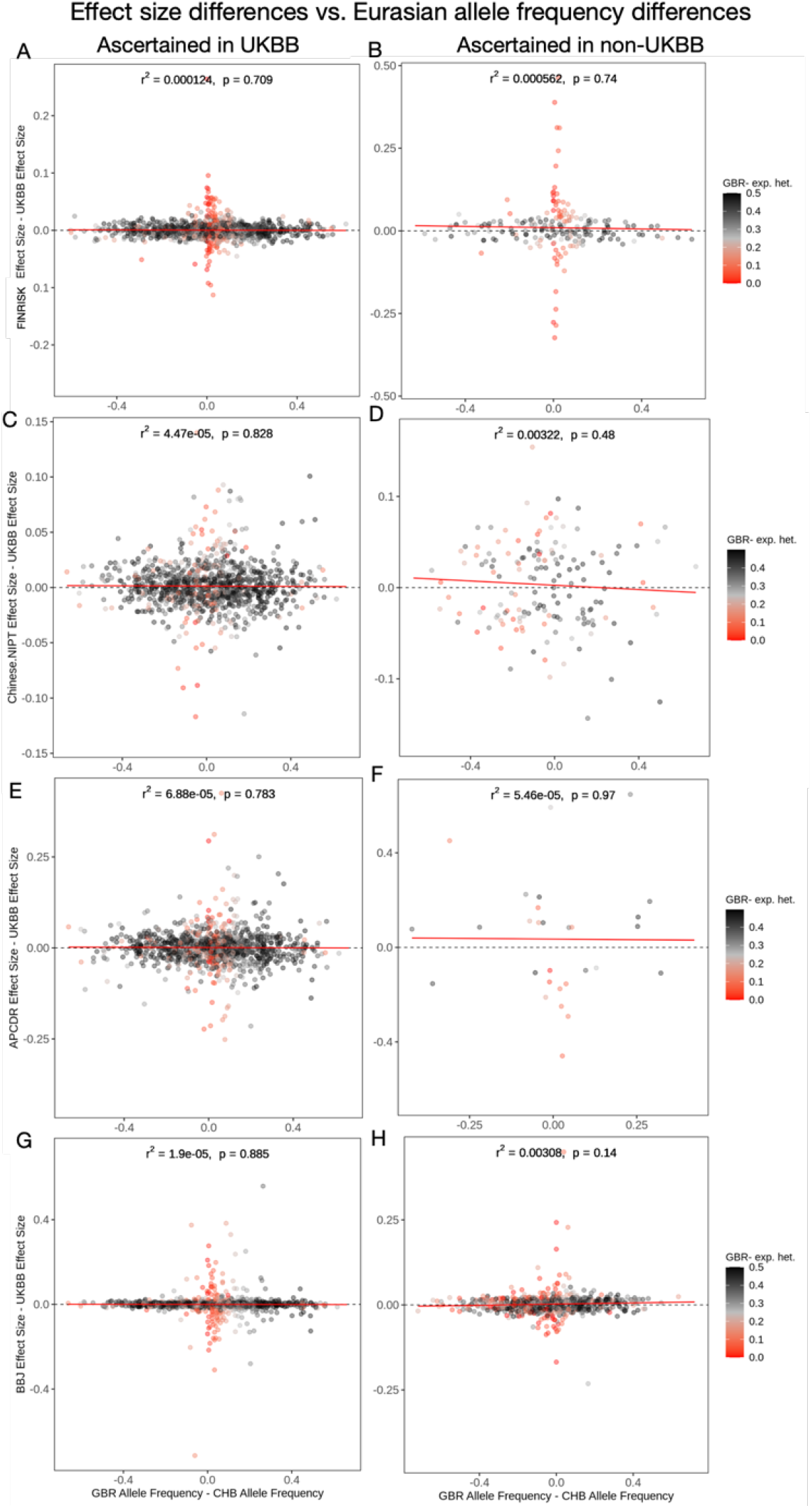
Each panel in this figure shows the same regressions as Figure S21, except that the difference in allele frequency is between the European and African population panels (GBR and LWK). SNPs are colored by their expected heterozygosity (2p(1-p)) in the GBR population.

**Figure S22.**
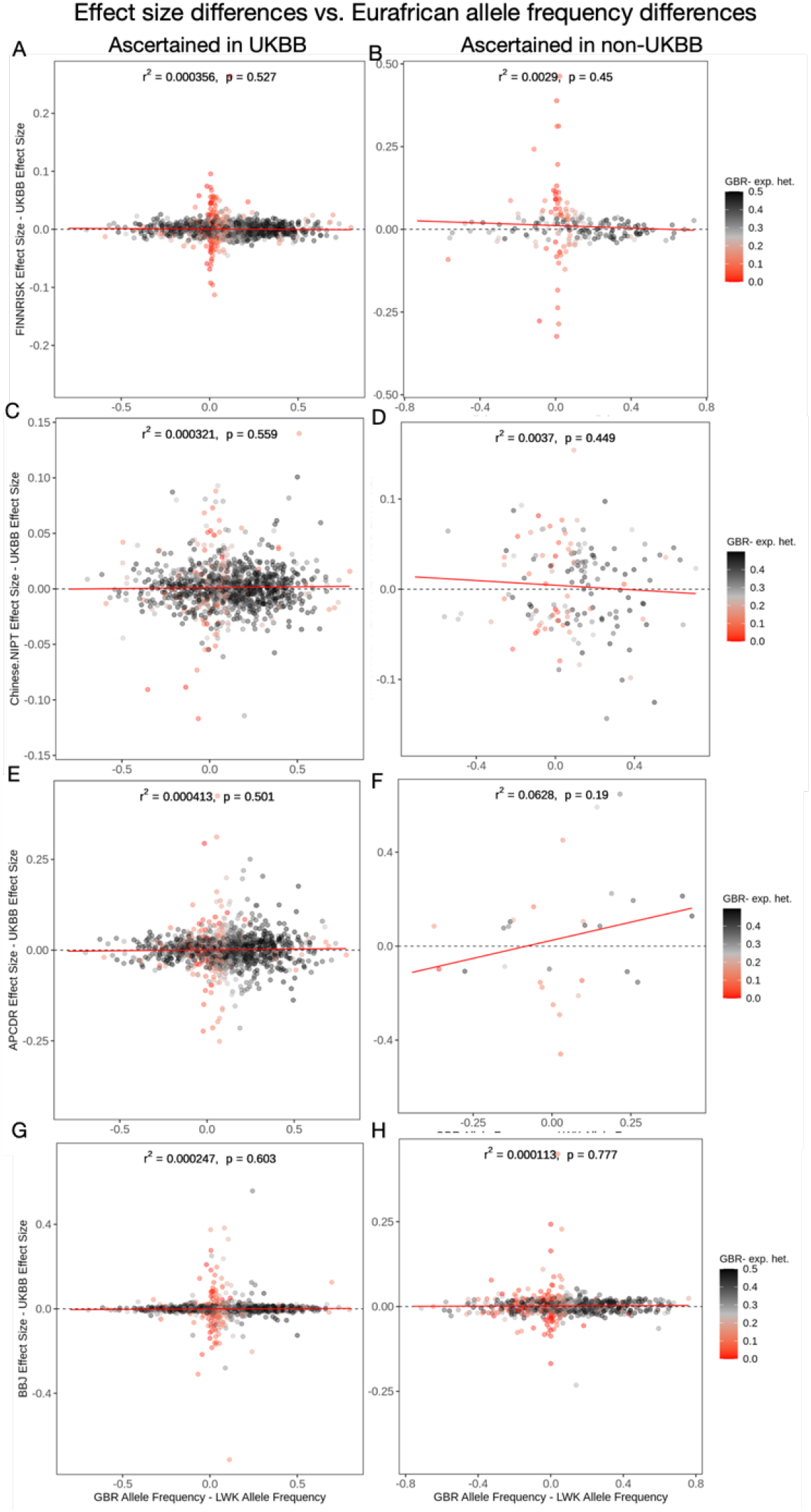
Each panel in this figure shows the same regressions as Figure S21, except that the difference in allele frequency is between the European and Asian population panels (GBR and CHB). SNPs are colored by their expected heterozygosity (2p(1-p)) in the GBR population.

**Figure S24.**
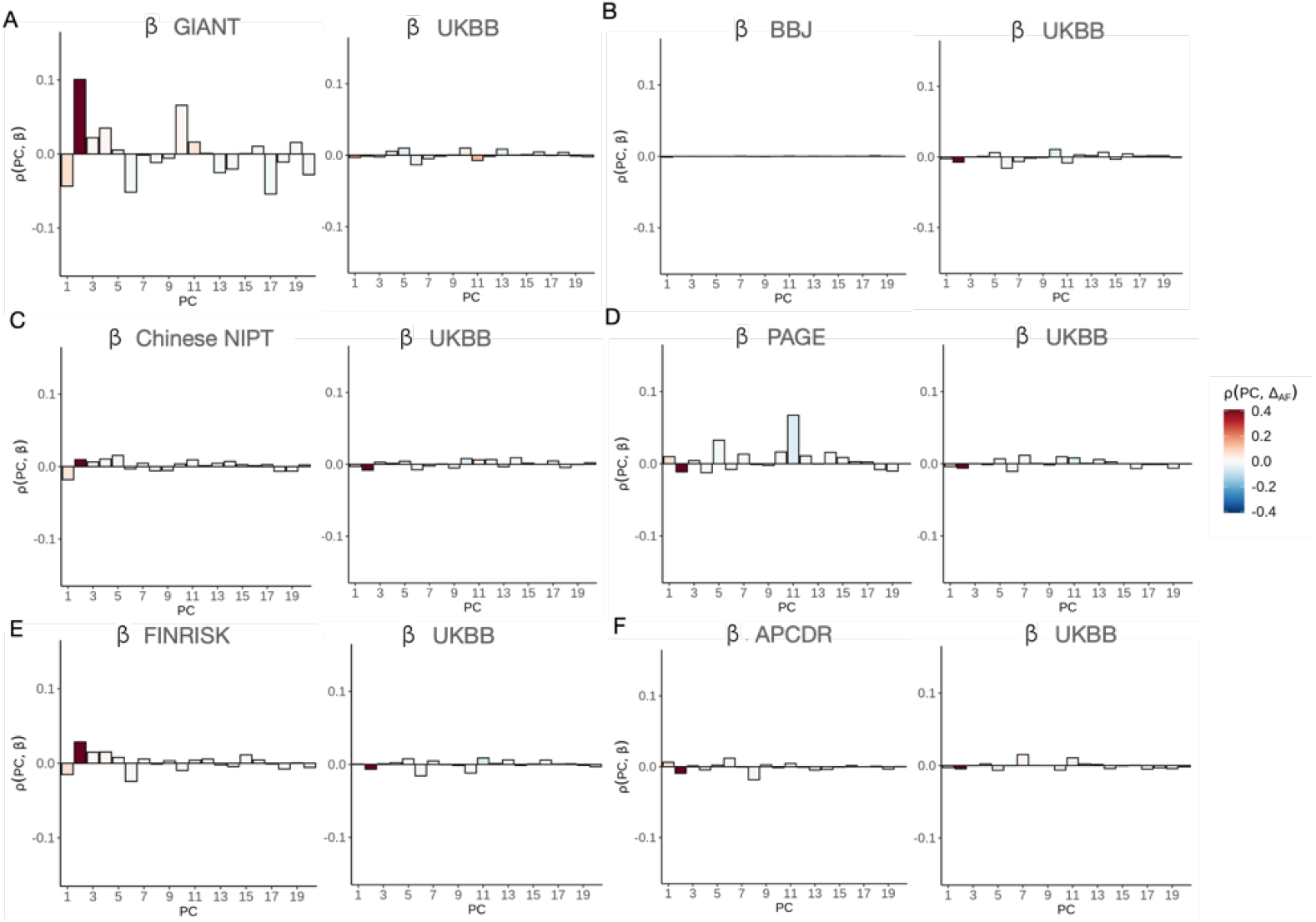
Pearson correlations between 20 PC loadings and height effect size estimates from a non-UKBB GWASs, compared to the same correlation using effect size estimates from the UKBB GWAS, for different choices of non-UKBB GWAS. The correlations were computed using SNPs that are present in both the UKBB and non-UKBB GWAS cohorts, and in the 1000 Genomes Project. The barplots are coloured by the correlation between each loading and the allele frequency difference between GBR and CHB. A) GIANT vs. UKBB. B) BBJ vs. UKBB. C) Chinese NIPT vs. UKBB. D) PAGE vs. UKBB. E) FINRISK vs. UKBB. F) APCDR vs. UKBB.

**Figure S25.**
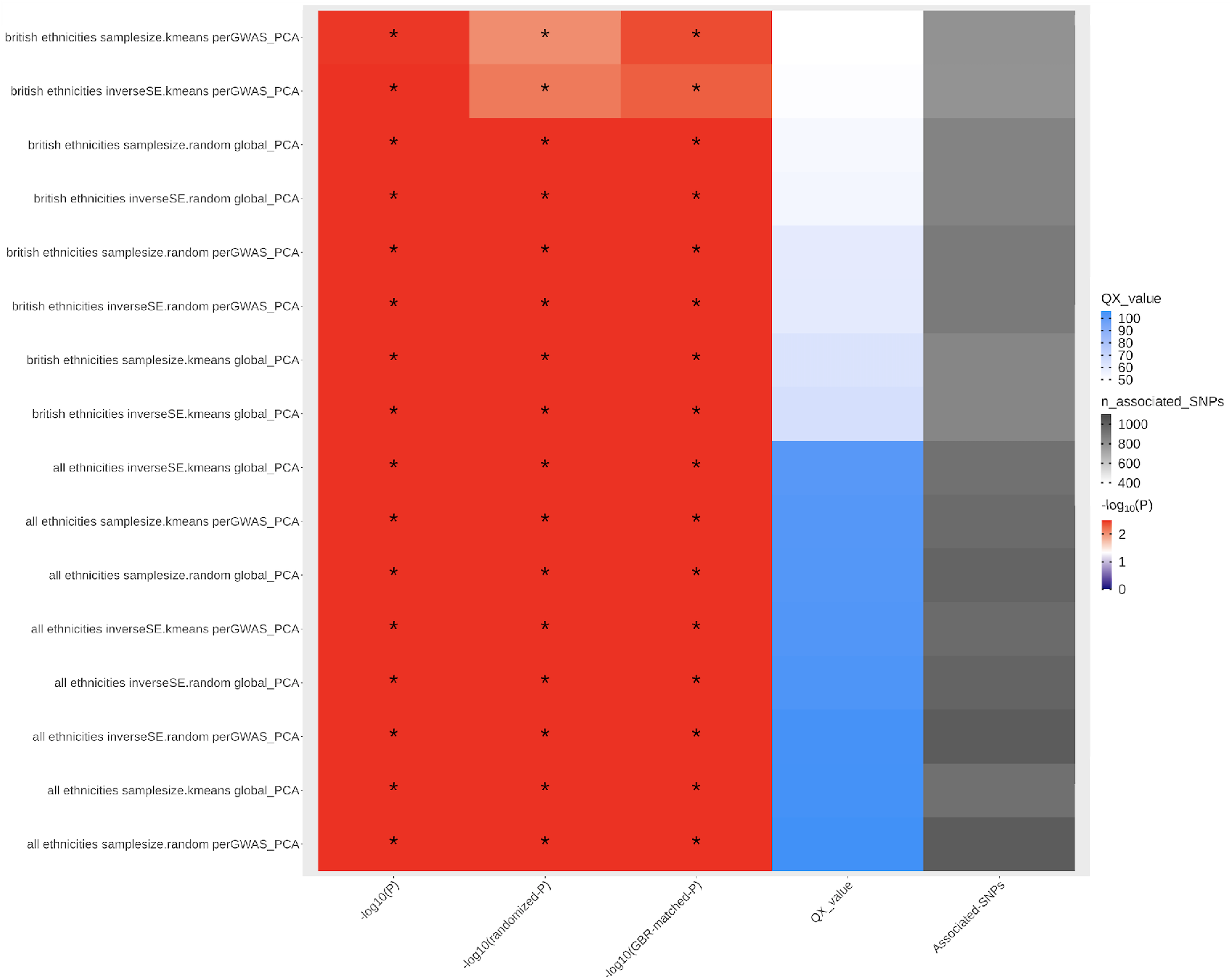
*Q*_*X*_ statistics and P-values for the trait height, obtained by using different types of meta-analysis methods on the UKBB data (inverse variance and sample size based) and two PCA correction approaches (global vs. per-GWAS). The meta-analyses were performed in “all ethnicities” as well as “White-British” set of individuals. The asterisk denotes a significance cutoff for *Q*_*X*_ of *P* < 0.05. “-log10(Chi squared P)” = -log10(P-value), obtained assuming a chisquared distribution for the *Q*_*X*_ statistic. “-log10(randomized P)” = -log10(P-value), obtained using the effect sign-randomization scheme. All other P-values were obtained by sampling random SNPs from the genome using the allele frequency matching scheme in different populations, as described in the Methods section.

**Figure S26.**
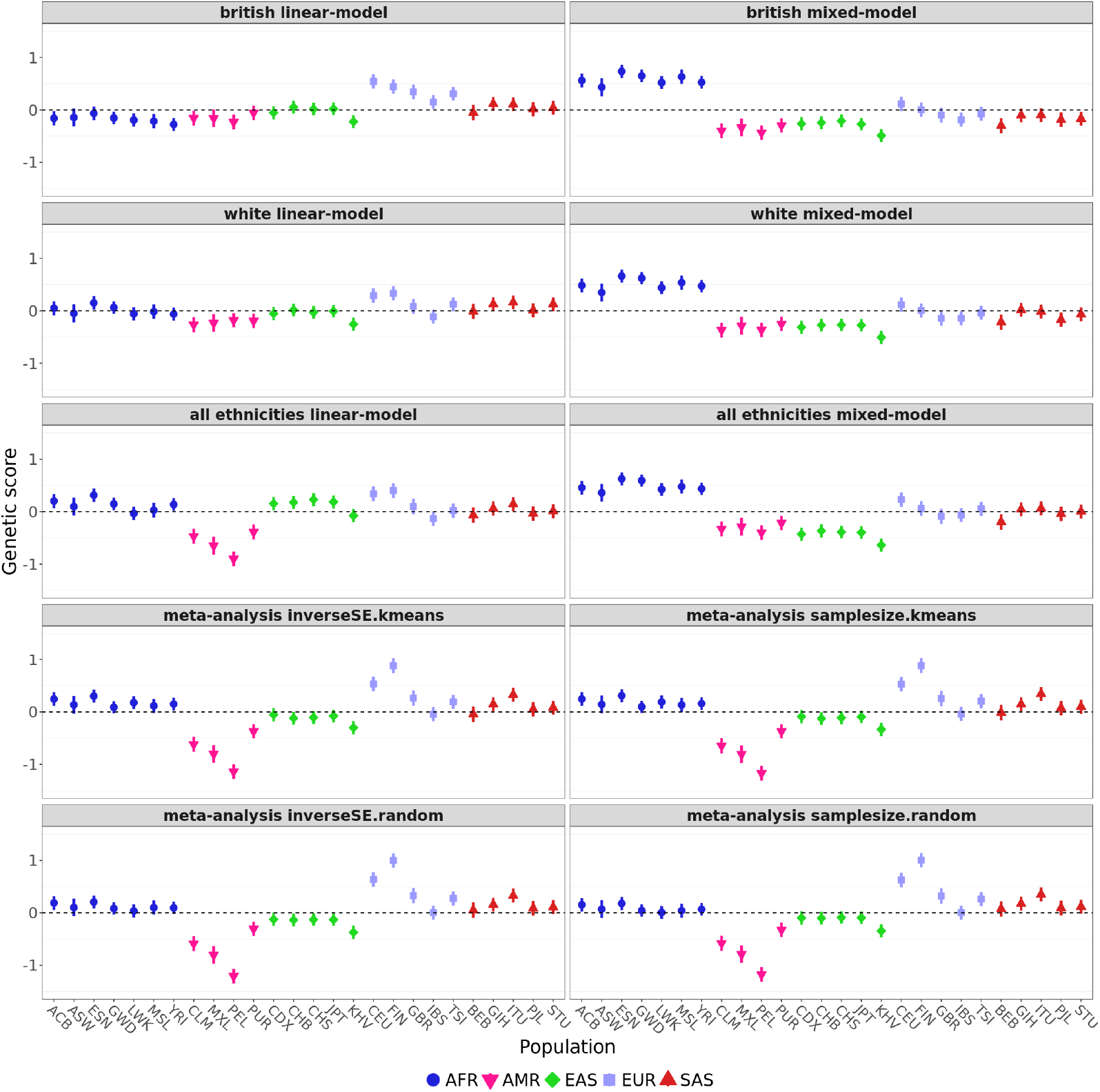
Polygenic scores for height in each of the 1000 Genomes population panels, using effect size estimates from UKBB, obtained via different types of meta-analysis methods (inverse variance and sample size based) and two PCA correction approaches (global vs. per-GWAS). The meta-analyses were performed in the “all ethnicities” as well as in the “White-British” set of UKBB individuals. Error bars denote 95% credible intervals, constructed using the method in Sohail et al., 2019, assuming that the posterior distribution of the underlying population allele frequency is independent across populations and SNPs.

**Figure S27.**
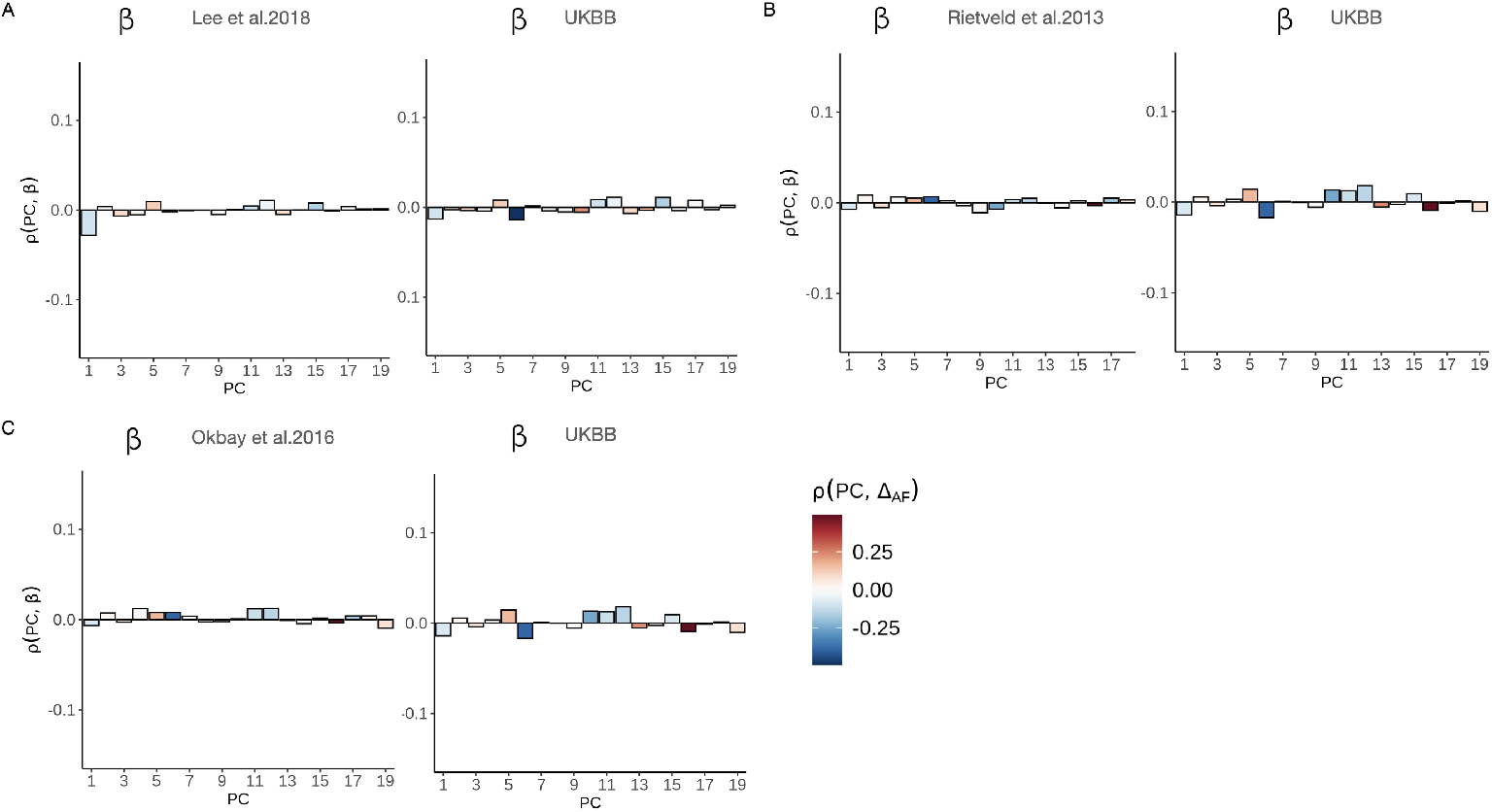
Pearson correlations between 20 PC loadings and educational attainment effect size estimates from a non-UKBB GWAS, compared to the same correlation using effect size estimates from the UKBB GWAS, for different choices of the non-UKBB GWAS. The correlations were computed using SNPs that are present in both the UKBB and non-UKBB GWAS cohorts, and in the 1000 Genomes Project. The barplots are coloured by the correlation between each loading and the allele frequency difference between GBR and TSI. A) Lee et al. 2016 vs. UKBB. B) Rietveld et al. 2018 vs. UKBB. C) Okbay et al. 2016 vs. UKBB.

## Supplementary Tables

**Table S1.** Full population descriptions of 1000 Genomes Project panels used in our analysis.

| Population code | Description | Super population |
| --- | --- | --- |
| CHB | Han Chinese in Beijing, China | EAS |
| JPT | Japanese in Tokyo, Japan | EAS |
| CHB | Southern Han Chinese | EAS |
| CDX | Chinese Dai in Xishuangbanna, China | EAS |
| KHV | Kinh in Ho Chi Minh City, Vietnam | EAS |
| CEU | Utah Residents with Northern and Western European Ancestries | EUR |
| TSI | Toscani in Italia | EUR |
| FIN | Finnish in Finland | EUR |
| GBR | British in England and Scotland | EUR |
| IBS | Iberian Population in Spain | EUR |
| YRI | Yoruba in Ibadan, Nigeria | AFR |
| LWK | Luhya in Webuye, Kenya | AFR |
| GWD | Gambian in Western Divisions in the Gambia | AFR |
| MSL | Mende in Sierra Leone | AFR |
| ESN | Esan in Nigeria | AFR |
| ASW | Americans of African Ancestry in SW USA | AFR |
| ACB | African Caribbeans in Barbados | AFR |
| MXL | Mexican Ancestry from Los Angeles USA | AMR |
| PUR | Puerto Ricans from, Puerto Rico | AMR |
| CLM | Colombians from Medellin, Colombia | AMR |
| PEL | Peruvians from Lima, Peru | AMR |
| GIB | Gujarati Indian from Houston, Texas | SAS |
| PJL | Punjabi from Lahore, Pakistan | SAS |
| BEB | Bengali from Bangladesh | SAS |
| STU | Sri Lankan Tamil from the UK | SAS |
| ITU | Indian Telugu from the UK | SAS |

**Table S2.** Full list of all traits tested and the total number of individuals (N) is shown for the GWAS in which this data was available. For GWAS summary statistics with variable N across positions, we list the maximum N for that study.

| Traits |  | UKBB | BBJ | FINRISK | PAGE | APCDR | Chinese<br>NIPT | GIANT |
| --- | --- | --- | --- | --- | --- | --- | --- | --- |
| Anthropometric | BMI | 359,983 | 158,284 | 24,722 | 49,335 | - | 60,652 | 231,410 |
|  | Height | 360,388 | 159,095 | 24,725 | 49,781 | 4,778 | 61,321 | 253,288 |
|  | Waist circumference | 360,564 | - | - | 33,942 | - | - | - |
|  | Waist hip ratio (WHR) | - | - | 24,671 | 33,904 | - | - | - |
| Blood pressure | diastolic blood pressure (DBP) | 340,162 | 136,615 | 21,588 | 35,433 | - | - | - |
|  | Systolic blood pressure (SBP) | 340,159 | 136,597 | 21,591 | 35,433 | - | - | - |
| Inflammatory | Platelet | 361,141 | 108,208 | 6,404 | 29,328 | - | - | - |
|  | Monocyte | 349,856 | 62,076 | - | - | - | - | - |
|  | C-reactive protein (CRP) | - | 75,391 | 16,529 | 28,537 | - | - | - |
|  | Hemoglobin (HbA1c) | 350,474 | 42,790 | - | 11,178 | - | - | - |
|  | Mean corpuscular hemoglobin concentration (MCHC) | 350,468 | 108,728 | - | 19,803 | - | - | - |
|  | Mean corpuscular hemoglobin (MCH) | 350,472 | 108,054 | - | - | - | - | - |
|  | Mean corpuscular volume (MCV) | 350,473 | 108,256 | - | - | - | - | - |
|  | Basophil count | 349,856 | 62,076 | - | - | - | - | - |
|  | Hematocrit count | 350,475 | 108,757 | - | - | - | - | - |
|  | White blood cell count (WBC) | 350,470 | 107,964 | - | 28,534 | - | - | - |
|  | Red blood cell count (RBC) | 350,475 | 108,794 | - | - | - | - | - |
|  | Lymphocyte count | 349,856 | 62,076 | - | - | - | - | - |
|  | Monocyte count | 349,856 | 62,076 | - | - | - | - | - |
|  | Neutrophil count | 349,856 | 62,076 | - | - | - | - | - |
| Kidney-related | Creatinine | 350,812 | 142,097 | 6,376 | - | - | - | - |
| Liver-related | Billirubin | - | 110,207 | - | - | 4,778 | - | - |
| Metabolic | Cholesterol | 361,141 | 128,305 | - | - | 4,778 | - | - |
|  | High density lipoprotein (HDL) | - | 70,657 | 21,620 | 33,063 | 4,778 | - | - |
|  | Low density lipoprotein (LDL) | - | 72,866 | 21,250 | 32,221 | 4,778 | - | - |
|  | Triglycerides | - | 105,597 | 21,619 | 33,096 | 4,778 | - | - |
|  | Glucose | - | 93,146 | 4,418 | 23,923 | - | - | - |
| Electrolits | Potassium | 350,053 | 132,938 | - | - | - | - | - |
|  | Sodium | 350,061 | 127,304 | - | - | - | - | - |
| Lifestyle | Cigarettes per day | 25,348 | - | - | 15,862 | - | - | - |

**Table S3.** Pairwise Fst computed using SNPs present in PAGE, to estimate population differentiation between PAGE and each of the 1000 Genomes Project panels.

| Population panel | Super-population | $F_{st}$ against PAGE |
| --- | --- | --- |
| PUR | AMR | 0.013 |
| CLM | AMR | 0.017 |
| MXL | AMR | 0.023 |
| ASW | AFR | 0.026 |
| PJL | SAS | 0.029 |
| BEB | SAS | 0.03 |
| GIH | SAS | 0.033 |
| ITU | SAS | 0.033 |
| STU | SAS | 0.033 |
| IBS | EUR | 0.034 |
| TSI | EUR | 0.035 |
| CEU | EUR | 0.036 |
| FIN | EUR | 0.036 |
| GBR | EUR | 0.036 |
| ACB | AFR | 0.045 |
| PEL | AMR | 0.054 |
| LWK | AFR | 0.058 |
| CHB | EAS | 0.062 |
| JPT | EAS | 0.062 |
| KHV | EAS | 0.062 |
| GWD | AFR | 0.063 |
| CHS | EAS | 0.064 |
| YRI | AFR | 0.064 |
| ESN | AFR | 0.065 |
| MSL | AFR | 0.065 |
| CDX | EAS | 0.066 |

**Table S4.**
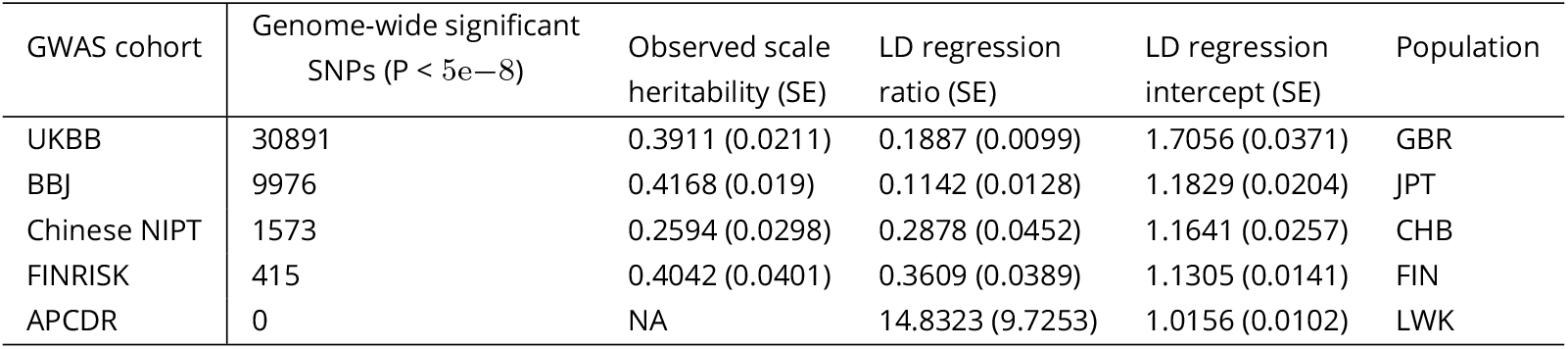
SNP-based heritability and LD Score regression ratio and intercept estimates (with standard errors in parentheses) for height measured in different cohorts. LD scores were computed using the closest population in the 1000 Genomes Project to each GWAS cohort (meta-analyses of multiple populations were not included here, but see Table 4). The APCDR heritability estimate is not shown because it was estimated to be negative, due to the small sample size of the cohort. For Chinese NIPT GWAS, we filtered out all the sites with INFO scores less than 0.4.

**Table S5.** Pairwise Pearson correlation coefficient between height effect size estimates from the UKBB GWAS and from another GWAS. The SNPs used were determined based on their P-value in the UKBB. n = number of SNPs used to compute the correlation.

|  | GIANT | FINRISK | PAGE | Chinese NIPT | BBJ | APCDR | SNP filtering scheme |
| --- | --- | --- | --- | --- | --- | --- | --- |
| Filtering based on UKBB P-values | 0.867<br>(n = 1703) | 0.531<br>(n = 1703) | 0.339<br>(n = 1703) | 0.510<br>(n = 1703) | 0.219<br>(n = 1703) | 0.061<br>(n = 1703) | SNP with lowest P-value in block |
| | 0.948<br>(n = 1059) | 0.738<br>(n = 1124) | 0.624<br>(n = 1139) | 0.625<br>(n = 1064) | 0.317<br>(n = 1097) | 0.107<br>(n = 1100) | SNP with lowest P-value in block, if $P < 1e-5$ |
| | 0.958<br>(n = 809) | 0.790<br>(n = 870) | 0.615<br>(n = 881) | 0.654<br>(n = 814) | 0.359<br>(n = 783) | 0.087<br>(n = 856) | SNP with lowest P-value in block, if $P < 5e-8$ |

**Table S6.** Pairwise Pearson correlation coefficient between height effect size estimates from the UKBB GWAS and from another GWAS. The SNPs used were determine based on their P-value in the non-UKBB study. n = number of SNPs used to compute the correlation.

|  | GIANT | FINRISK | PAGE | Chinese NIPT | BBJ | APCDR | SNP filtering scheme |
| --- | --- | --- | --- | --- | --- | --- | --- |
| Filtering based on non-UKBB P-values | 0.698<br>(n = 1703) | 0.167<br>(n = 1703) | 0.128<br>(n = 1703) | 0.335<br>(n = 1703) | 0.018<br>(n = 1703) | 0.121<br>(n = 1703) | SNP with lowest P-value in block |
| | 0.931<br>(n = 738) | 0.322<br>(n = 199) | 0.258<br>(n = 366) | 0.687<br>(n = 157) | 0.561<br>(n = 709) | 0.138<br>(n = 29) | SNP with lowest P-value in block if $P < 1e-5$ |
| | 0.938<br>(n = 570) | 0.334<br>(n = 136) | 0.260<br>(n = 257) | 0.654<br>(n = 112) | 0.732<br>(n = 412) | 0.087<br>(n = 15) | SNP with lowest P-value in block if $P < 5e-8$ |

**Table S7.** Height *Q*_*X*_ scores when using LD blocks derived from closely related populations. In the case of BBJ and the Chinese biobank, we used the ASN-specific LD blocks. In the case of APCDR, we used the AFR-specific blocks. In the case of PAGE, we used the AFR-specific blocks. The left columns show scores obtained when we used a *P* < 1_e_−5 threshold to include SNPs in the polygenic scores. The right columns show scores obtained using the genome-wide significant threshold (*P* < 5_e_−8). The number of trait-associated SNPs used to compute the scores are shown for both cutoffs.

| GWAS cohort | Lenient P-value threshold ($1e-5$) | | Strict P-value threshold ($5e-8$) | |
| --- | --- | --- | --- | --- |
|  | Num. trait-associated SNPs | Qx | Num. trait-associated SNPs | Qx |
| BBJ | 693 | 9.701 ($P = 0.99$) | 401 | 36.00 ($P = 9e-2$) |
| Chinese NIPT | 160 | 13.05 ($P = 0.98$) | 59 | 26.41 ($P = 0.44$) |
| PAGE | 592 | 97.23 ($P = 4e-10$) | 94 | 65.86 ($P = 3e-5$) |
| APCDR | 60 | 21.15 ($P = 0.73$) | 0 | NA |

## Notes

### Competing Interest Statement

The authors have declared no competing interest.

### Summary of Updates

Version 5 of this preprint has been peer-reviewed and recommended by Peer Community In Evolutionary Biology (https://doi.org/10.24072/pci.evolbiol.100125).

